# Stiffening cells with light

**DOI:** 10.1101/2025.01.30.635790

**Authors:** Eva Gonzalez, Jana El Husseiny, Finn Bastian Molzahn, Tiffany Campion, Hadrien Jalaber, Stéphanie Dogniaux, Pierre-Henri Puech, Oliver Nüsse, Laure Gibot, Julien Husson

## Abstract

Fluorescence microscopy is widely used to observe structures and dynamic processes in living cells and organisms and is often used as if it were purely innocuous to the cells or structures of interest. However, it can lead to phototoxicity, which can affect the cellular behavior and lead to erroneous interpretations of the observations. The major cause of cell damage through phototoxicity is the production of reactive oxygen species (ROS), which can form crosslinks between intracellular molecules, including proteins and nucleic acids. By using profile microindentation and atomic force microscopy, we demonstrate that the excitation of various fluorescent probes leads to a large increase in the stiffness of several cell types within seconds of illumination. The stiffening exhibits a dose-dependent response, where longer exposure times to exciting light are correlated with larger stiffening.

This photostiffening effect explains why T cells loaded with the Fluo-4 Calcium probe stop emitting a protrusion within seconds after the excitation light is turned on. We observed photostiffening in different cell types and fluorophores. We showed that repeated cell indentation alone led to cell stiffening as well as excitation with blue or UV light in the absence of a fluorophore. However, in the latter case, the stiffening was much smaller than that when the fluorophore was excited. We used both sharp and blunt indenters to show that stiffening occurred not only at the cell cortex level but also deeper in the cell interior and that photostiffening is independent of actin cytoskeleton organization. We correlated the increase in cell stiffness with the production of intracellular reactive oxygen species and reproduced cell stiffening by incubating cells with a ROS inducer, H_2_O_2_. The excitation of the photosensitizer Pheophorbide *a*, which induces a specific type of reactive oxygen species, namely singlet oxygen, also led to cell stiffening. This study reminds the experimentalists that it is crucial to perform controls when using fluorescence. It further allows us to propose exploiting photostiffening as a new method for rapidly quantifying phototoxicity.

**Significance:** This study reveals a direct relationship between fluorescence excitation and cell mechanical properties. The generality of th is phenomenon across diverse fluorophores and cell types highlights the importance of controlling phototoxicity in fluorescence experiments, in particular in complex, quantitative cell biology and biophysical experiments, but also reveals an unexpected rol e of intracellular reactive oxygen species production and identifies cell stiffness as a proxy for assessing the efficacy of photodynamic therapy.

## 1. Introduction

Fluorescence microscopy is widely used to observe structures and dynamic processes in living cells and organisms because it allows selective and specific imaging of selected molecules. However, fluorescence microscopy is not innocuous and may lead to phototoxicity, which can affect cellular behavior and lead to erroneous interpretations of observations. The major cause of cell damage through phototoxicity is the local production of reactive oxygen species (ROS) [Icha, 2017].

ROS are chemically reactive molecules that are produced naturally within cells as a byproduct of metabolism or by specific enzymes called NADPH oxidases. They are essential components of cell signaling and regulate many physiological processes [Sies, 2022]. Nevertheless, ROS overproduction can be harmful up to inducing cell death. ROS are produced by light through fluorophores or endogenous photosensitizers such as flavins and porphyrins, which absorb light and react with oxygen [Icha, 2017]. Furthermore, most of the time, minutes-to-hours-long living cell microscopy experiments are performed with an atmospheric concentration of oxygen i.e. 20%, which is much larger than the physiological tissue oxygen concentration of about 4-5%, favoring ROS overproduction during experiments. A consequence of ROS overproduction is the oxidation of DNA, proteins, and lipids. ROS can also form crosslinks between proteins [Feng, 2022], including covalent bonds through cysteines [Ahmad, 2017], and crosslinks and between DNA and proteins [Olinski, 1992]. Most fluorescence users are aware of the existence of phototoxicity and attempt to reduce it empirically. For instance, in reconstituted cytoskeleton systems such as microtubules in the absence of oxygen scavengers in the surrounding medium, fluorescent microtubules break down quickly under fluorescence excitation. When studying cells, researchers usually perform a phototoxicity assay to assess the close to no bias status of their imaging techniques. Monitoring the cell cycle (entry into mitosis, proliferation) has been widely used [Douthwright, 2017; Laissue, 2017], but it also has limitations at low light intensity [Alghamdi, 2021]. Thus, other parameters have been used to assess phototoxicity, such as cell mobility [Alghamdi, 2021] and cell protrusion rates [Mubaid, 2017]. These tests assess the long-term effects of light but not its possible immediate effects.

In preliminary studies, we worked with T cells loaded with the calcium probe Fluo-4, which apparently inhibited the deformation of the cells during their activation by antibody-covered microbeads: T cells appeared “frozen” when the fluorophore was excited. This artifact was one of the initial hints that led us to ask whether the fluorescence excitation of a fluorophore could influence the capacity of cells to deform and if it could make them stiffer. This might seem counterintuitive, as one could think that potential phototoxicity would break the cytoskeleton, which is commonly thought to provide mechanical properties to cells [Galie, 2022]. Thus, phototoxicity could be expected to lead to cell softening. Consistent with this, some studies have shown that ROS can disrupt the actin cytoskeleton [Milzani, 1997; Brival, 2024], and others have observed a decrease in cell stiffness due to phototoxicity.

In this study, we first demonstrated the effect of Fluo-4 excitation on T-cell stiffness and its capacity to deform during activation. We then asked whether we could expand our observations to different fluorophores and cell types. To achieve this, we used an innovative profile microindentation technique that we have developed and refined [Husson, 2023]. As we show here, we obtained a totally opposite result, as expected from the above-mentioned literature, with a huge stiffening of the cells. We showed that this stiffening is correlated with intracellular ROS production, that ROS generation independent of light induces cell stiffening, and that photosensitizer excitation also induces cell stiffening. A positive control was made by incubating both T cells and Bovine aortic endothelial cells (BAECs) with H_2_O_2_, and in both cases, the cells stiffened. We extended this positive result to pheophorbide *a*, which is classically used as a photosensitizer in photodynamic therapy (PDT) and specifically induces singlet oxygen [Sindhu, 2018].

## 2. Results

### 2.1 Fluo-4 excitation leads to irreversible T-cell deformation

Given that Fluo-4 excitation has been observed to stop cell protrusions and limit cell migration, we tested Fluo-4 excitation and asked whether this could more generally change the way cells deform. Thus, we wanted to monitor whether their mechanical properties would change. To do so, we loaded T cells with Fluo-4 using conventional protocols, pressed micropipette-held cells against a rigid glass wall to deform them, turned on blue light to excite Fluo-4 fluorescence, and observed if, under fluorescence excitation, T cells could recover their initial shape (Figure 1A,B). In the absence of fluorescence excitation, control cells recovered their initial shape over a typical timescale of 5-10 seconds (Figure 1C). However, under excitation light, cells did not recover at all: these cells were totally “frozen” (or “cooked”, and as such we informally called them “cell pancakes”). If any, the shape recovery was so slow that we could not measure it, i.e. the timescale for recovery was too large. We present an example of these experiments in Supplementary Movie S1.

**Figure 1.**
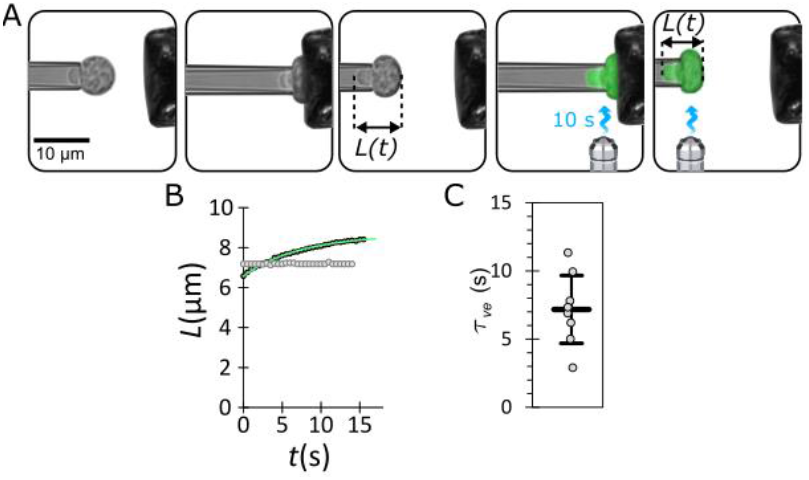
(A) Fluo-4 excitation leads to irreversible T-cell deformation. T cells loaded with Fluo-4 were held using a micropipette and pressed against a rigid wall for a few seconds. The micropipette was rapidly retracted, and the cell length L(t) was recorded over time (first three images on the left). After typically 20 s, the cell was pressed a second time against the rigid wall, but while deformed, a blue fluorescence excitation light was turned on (at 100% power) and left on (two last images on the right, see Movie S1). (B) Example of cell length vs. time for the first cell recovery in the absence of excitation light (black dots) and the second recovery in the presence of excitation light on (gray dots). In the latter case, the cell barely deforms over time. In the first recovery, exponential saturation 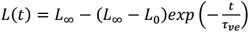 is fitted to the data, where *L*_0_ and *L*_∞_represent the initial and final cell lengths, respectively, and τ_*ve*_ is a viscoelastic recovery time. The fit is shown as a green solid line. (C) Viscoelastic recovery time measured during the first recovery (in absence of excitation light, for 3 donors and 8 cells). For illuminated cells, this time is so large that it cannot be fitted or estimated from our finite-duration experiments.

These observations led us to speculate that exciting Fluo-4 fluorescence would perturb the formation of protrusions by T cells during activation, a process that we have extensively described in previous work [Sawicka2017; Zak2021; Zucchetti2021]. Furthermore, as our previous experiments on “cell pancakes” were conducted using maximal lamp power, we asked whether using a lower lamp power, more typical for cell biology experiments, would translate into less perturbing cells, i.e., showing a dose-effect response.

### 2.2 Photostiffening affects T-cell protrusion during activation in a dose-dependent manner

We further asked whether photostiffening could influence cell active deformation during T cell activation. We activated CD4+ T cells with antibody-coated microbeads (see Material and Methods section). T cells were loaded with Fluo-4 calcium probe. In the absence of fluorescence excitation, T cell activation proceeded efficiently, as evidenced by the emission of a large protrusion, a process that we have previously comprehensively characterized [Sawicka, 2017; Zak, 2021; Zucchetti, 2021]. Interestingly, very quickly after blue light was applied to activating T cells, the protrusion stopped growing abruptly and cells stopped their typical morphological changes during activation (Figure 2). When we decreased the power levels of the blue lamp from 100% to 10%, the duration needed to stop the protrusion increased from 1.6 ± 0.3 s (4 donors, 16 cells, average ± SD) to 4.7 ± 1.2 s (3 donors, 9 cells, average ± SD).

**Figure 2.**
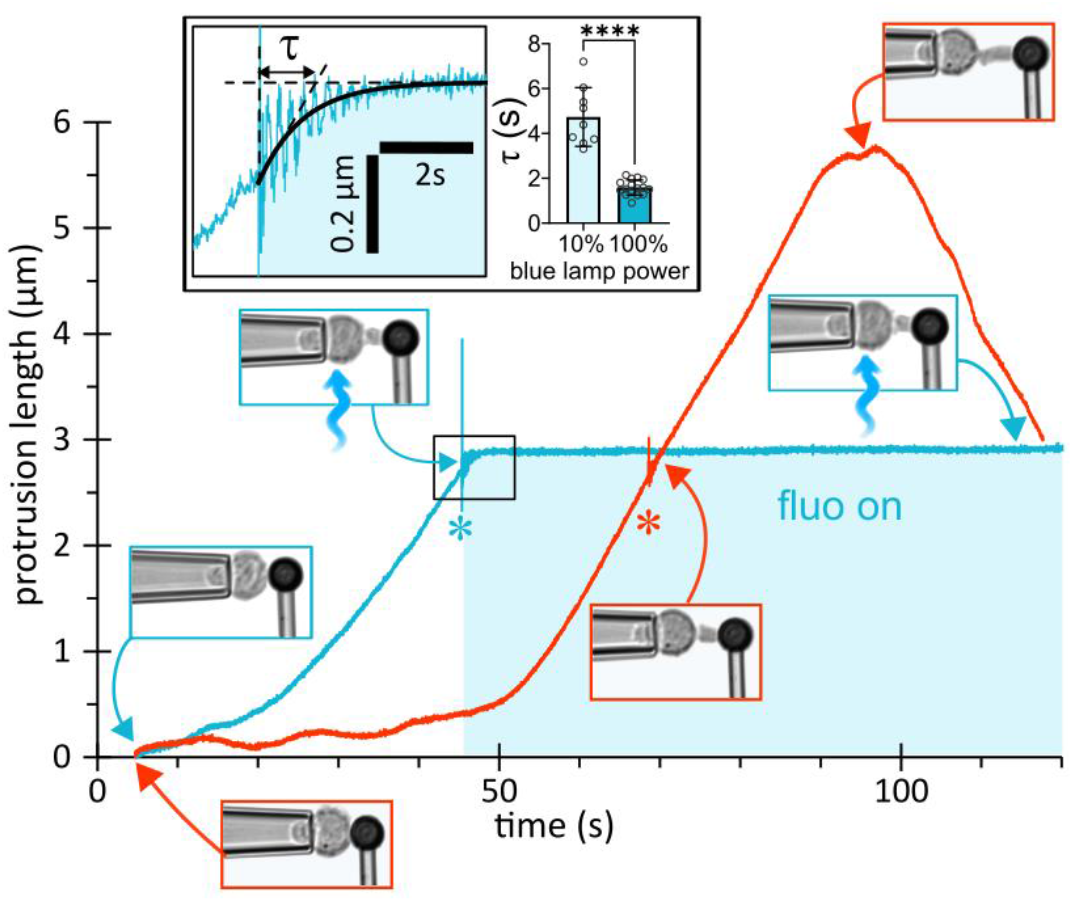
T-cell activation stops when Fluo-4 is excited. Fluo-4-loaded CD4 T cells were activated with anti-CD3+anti-CD28 -coated microbeads. In the absence of fluorescence excitation (red curve), T cells emit a large protrusion. Separately, in other cells, blue light was turned on (blue lamp power 100%) while the protrusion was growing, and very quickly after light application, the protrusion stopped growing (solid blue line, star symbol shown when excitation is turned on) within a timescale τ that depended on the blue lamp power (histogram in inset, right). The moment at which the excitation light is turned on is shown by inducing a small vibration into the cell-holding micropipette by gently flicking the micropipette holder (see inset, left). In the control case (no excitation, red curve), the flicking was also performed to exclude any potential perturbation due to the vibration. An example is shown in Movie S2.

### 2.3 Microindentation allows quantifying mechanical changes in cells exposed to excitation light

We used profile microindentation to quantify cell stiffness while exposing the cells to fluorescence excitation. The setup consists of holding a non-adherent (or detached) cell with a micropipette, pushing it against a flexible micropipette ending with a glass sphere and used as a cantilever (Figure 3). The setup is mounted on an epifluorescence inverted microscope, and indentations can be alternated with fluorescent light exposure. One indentation will be called a “cycle” hereafter. We implemented different protocols during which cells were exposed to a single 10-s flash of excitation light of given intensity or several consecutive 4-s pulses. We also used a two-color protocol in which one fluorophore was used to induce cell stiffening and the other to determine intracellular ROS production (Figure 3). Unless stated otherwise, micro-indentation experiments were performed using a 100X oil immersion objective. However, we performed some control experiments using a 40x air objective to verify the impact of the optic configuration on the perturbations of the cells (see Figure 4E-F and Figure 6).

**Figure 3.**
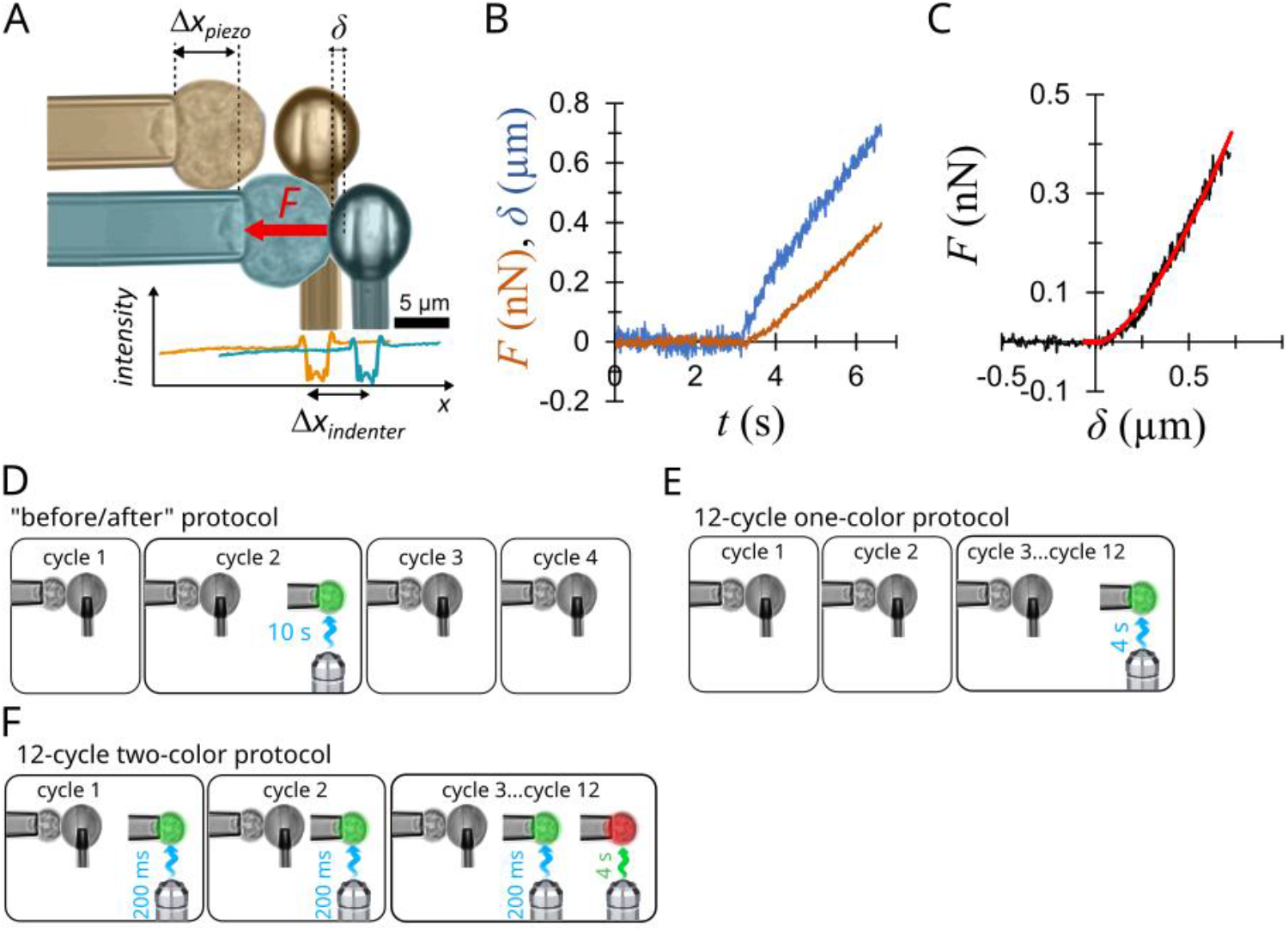
**(A)** Profile microindentation using a spherical microindenter [Husson, 2023; Guillou, 2016]. A stiff micropipette holds a cell (left) and is placed close to the tip of a microindenter consisting of a flexible micropipette ending with a glass bead (right). Translation of a single-axis translation stage controlled by a piezo actuator along the horizontal *x*-axis (*Δx*_*piezo*_) leads to the compression of the cell against the microindenter (blue), resulting in the indentation of the cell of amplitude *δ*. The deflection of the microindenter is equal to the translation of the tip of the microindenter, *Δx*_*indenter*_. The force *F* exerted by the microindenter on the cell is *F* = *k*_*indenter*_*Δx*_*indenter*_, where *k*_*indenter*_ is the bending stiffness of the microindenter, which is determined independently. The indentation of the cell, *δ*, is deduced from the difference between the piezoelectric stage translation and the indenter’s deflection: *δ*=*Δx*_*piezo*_*-Δx*_*indenter*_. To accurately measure the position of the tip of the microindenter, the intensity profile of transmitted light (bright-field illumination) along the *x*-axis was analyzed at an acquisition frequency of approximately 400 Hz and correlated to a reference intensity profile acquired at the beginning of the experiment, leading to a detection accuracy of approximately 20 nm using a 100x magnification objective [Husson, 2023]. **(B)** Time traces acquired during the experiment: force (orange) and indentation (blue). Cell compression is performed by translating the piezoelectric stage at constant velocity (*dx*_*piezo*_/*dt* ranging from 0.25 to 0.5 µm/s) until a preset force threshold *F*_*max*_ is reached (for T cells and PLB cells *F*_*max*_ = 0.4 nN, and for BAECs *F*_*max*_ = 3 nN), after which the piezoelectric stage is withdrawn at a constant velocity, with no waiting time at contact. **(C)** Force-indentation curve (black curve). Data are fitted with the Hertz model 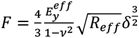 [Guillou, 2016, Husson, 2023, Markova, 2024] (red curve), where 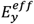 is the effective cell stiffness (called “cell stiffness” herein),νis the Poisson’s ratio taken as 0.5, *δ* is the indentation of the cell, and *R*_*eff*_ is an effective radius defined by 1/*R*_*eff*_ = 1/*R*_*indenter*_ + 1/*R*_*cell*_, where *R*_*indenter*_ is the radius of the microindenter tip or micropipette bead, and *R*_*cell*_ is the cell radius. **(D)** In the so-called “before-after” protocol, cells are first indented twice (cycles 1-2), and at the end of cycle 2, they are exposed to 10 s of excitation light. After this, cells are indented twice to monitor the eventual change in stiffness (cycles 3-4). Depending on the fluorophore used, the excitation light is either ultraviolet (UV, when using Hoechst, Pheophorbide *a* or encapsulated Pheophorbide *a*) or blue (when using Fluo-4 and of 2’,7’-dichlorodihydrofluorescein diacetate (DCFH-DA)). In some experiments using DCFH-DA, 200-ms snapshots of blue light were added in the sequence to monitor the level of DCFH-DA fluorescence used as a reporter of the ROS level. **(E)** In the so-called “12-cycle one-color protocol”, the cell is first indented twice without fluorescence excitation, and during cycles 3–12, it is indented and then exposed to 4 s of excitation light at each iteration (blue light if cells are loaded with Fluo-4). **(F)** In the so-called “12-cycle two-color protocol”, during the first two cycles, the cell is indented and exposed to a 200-ms snapshot of blue light to monitor the level of DCFH-DA fluorescence. Then, during cycles 3 to 12, the cell is indented, exposed to 200 ms of blue light, and then to 4 s of green light to excite the CellTracker Red probe at each iteration.

**Figure 4.**
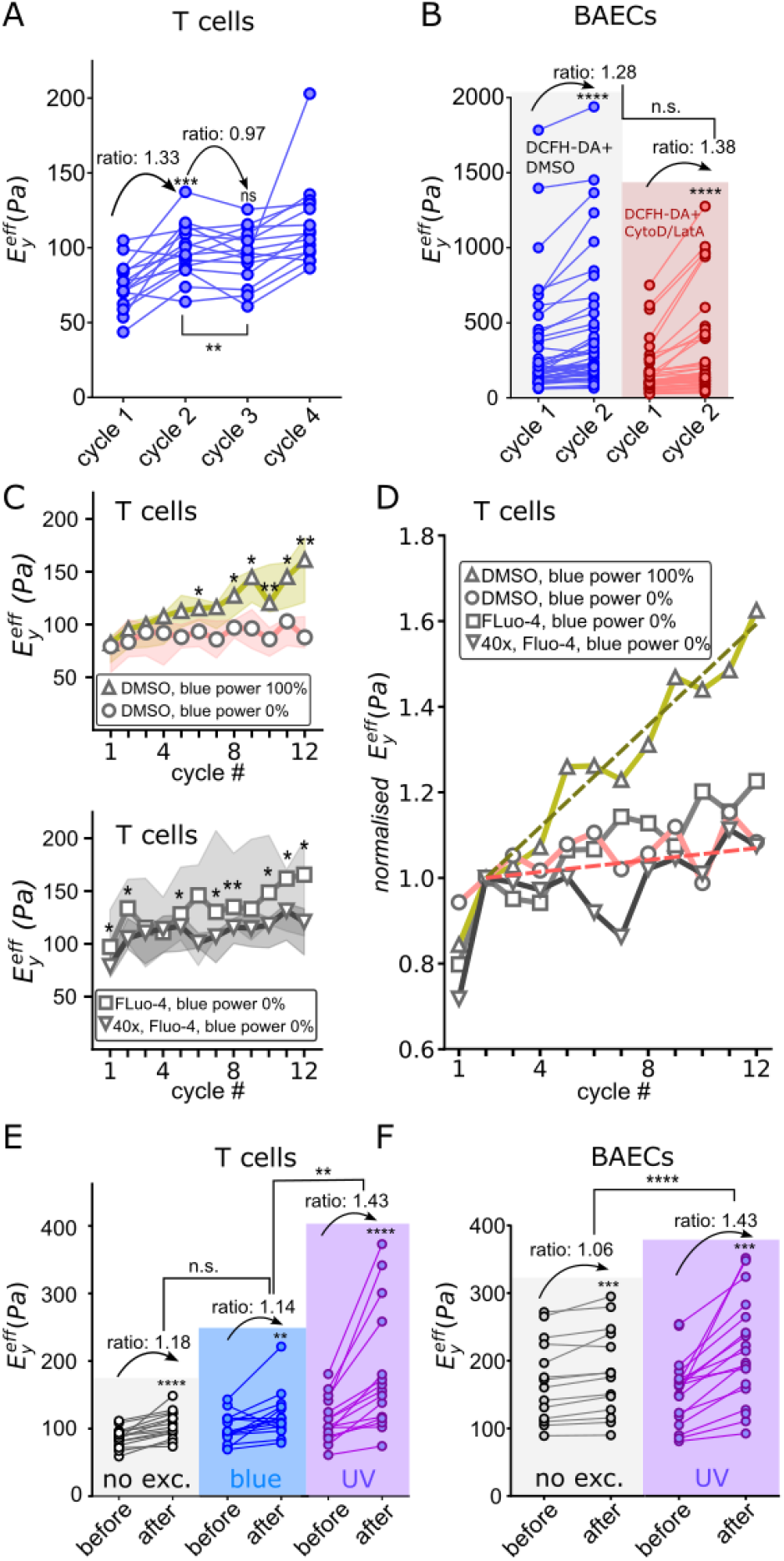
**(A)** Repeated indentation in the absence of excitation light induces cell stiffening, and a 10-s rest mitigates this increase. As an example, T cells were incubated with DMSO (see statistics in Supp. Table 1 for several fluorophores and cell types), th e median ratio between stiffness measured during the first and second cycle is 1.33, i.e., a 33% increase, while the stiffness does not significantly vary from cycle 2 to cycle 3 when cells are left resting 10-s without light excitation in-between indentation (median ratio from cycle 2 to 3: 0.97). **(B)** Inhibiting the actin cytoskeleton using Cytochalasin D together with Latrunculin A on BAECs preloaded with DCFH-DA does not avoid cell stiffening between cycle 1 and cycle 2 (i.e. before any light excitation). As expected, the cytoskeletal inhibitors led to a significantly lower initial stiffness, as measured during cycle 1 (a median of 111 Pa, n=46 cells, *vs*. a median of 171 Pa, n=44 cells). However, the increase in stiffness from cycle 1 to cycle 2 was not influenced by the presence of Cytochalasin D and Latrunculin A inhibitor (the median values of 28% vs 38% relative increase are not significantly different). **(C)** On top: T cells incubated in 0.2 % DMSO were subjected to the 12-cycle one-color protocol in the presence (blue power 100%) or absence of blue light excitation (blue power 0%). Light was applied starting at cycle 3, and cell stiffness in the presence of blue light became significantly larger for all cycles starting from cycle 8 on (Mann-Whitney test, n=9 cells). On the bottom: T cells were preloaded with Fluo-4 and submitted to the 12-cycle one-color protocol with no excitation light. Instead of the usual 100x-magnification (oil) objective, a 40-X (air) magnification objective was used. Dots are median values, and error bars are interquartile ranges. **(D)** Relative increase in stiffness: the data in panel C are normalized by the stiffness measured at cycle 2. The effect of blue light, in the absence of a fluorophore, can be seen with a relative increase of 5.9% increase per cycle versus a 1.2%, 1.1%, and 2.6% increase per cycle for DMSO without light, Fluo-4 without light and 40x magnification, and Fluo-4 without light, respectively. **(E)** Light alone has a stiffening effect: T cells were preincubated with DMSO and then submitted to the before-after protocol (Fig. 3), where they were kept either in the dark (left column), or exposed to blue (middle column) or UV light (right column) for 10 s. The increase resulting from blue light (14%) was not significantly different from that obtained in the absence of light excitation (18%), whereas UV light induced a significantly larger (43%) stiffness increase. **(F)** UV light in the absence of dye also stiffens endothelial cells: BAECs were not incubated with any fluorophore and submitted to the before-after protocol where they were either kept in the dark (left column) or exposed to UV light (right column) for 10 s. The stiffening in the presence of UV light (43%) was significantly greater than that in the absence of light excitation (6%).

### 2.4 Control experiments: effects of indentation and excitation light on cell stiffness

In this study, we addressed the effect of fluorophore excitation on cell stiffness using the so-called “before-after” protocol (Fig. 3), where each cell was first indented twice to measure its stiffness twice (cycles 1-2), then the cell was exposed to 10 s of excitation light, following which cell stiffness was measured twice again (cycles 3-4, see Figure 1). Before considering the effect of fluorophore excitation using this protocol, we asked whether (i) indentation itself would lead to cell stiffening and if (ii) letting cells rest for several seconds in between cycles, without indentation or excitation light would limit this stiffness increase. We observed that when indented for the second time (still before light exposure), the cell was 11–52% stiffer than during the first indentation: in Figure 4A we show the stiffness measured during cycles 1-4 for T cells incubated in 0.2% DMSO for control experiments. Supp. Table 1 presents the statistics for different fluorophores and cell types (T cells, BAECs, and PLB cells). Under all these conditions, this indentation-induced stiffening was present to various extents, in a cell-dependent manner, and as it occurred before light excitation, it was more likely a consequence of the indentation and not the loading with a fluorophore. Furthermore, we excluded the role of the change in cell geometry after the first indentation by examining the side images provided by our technique, i.e., the stiffening appears to be an intrinsic change in the mechanical properties.

We then asked whether the increase between the first and second indentation (i.e. between cycle 1 and cycle 2) was a mechanosensitive response of the actomyosin cortex. We ran the before-after protocol on BAECs with destabilized actomyosin cortex due to the simultaneous effects of Cytochalasin D and Latrunculin A. As expected, in the presence of these inhibitors, the stiffness measured during cycle 1 was lower than that of control cells incubated in medium in the absence of inhibitor (Fig. 4B). However, the increase from cycle 1 to cycle 2 still existed in the presence of the inhibitors (28% in absence of inhibitor, n=44 cells, 38% in presence of inhibitor, n=46 cells, p = 0.117, Mann-Whitney test). This suggests that the increase in stiffness between the first and second indentation is not an active actin-dependent response of the cell but rather a passive, possibly viscoelastic or plastic response.

We next asked whether adding a resting duration of 10 s in the dark in-between two indentations would avoid this stiffening effect, and indeed, for the three cell types and several fluorophores that we tested (Supp. Table 2), this rest led to at most a 6% increase in BAEC stiffness or no increase in stiffness for CD4 and PLB cells (see an example of T cells in Fig. 4A and the fourth column in Supplementary Table 2).

Next, we asked whether cell stiffness could be influenced by light excitation in the absence of fluorophore loading. We pre-incubated CD4 T cells with the fluorophore solvent dimethyl sulfoxide (DMSO) and submitted them to the before-after protocol, exposing cells to no light, blue light (455 nm, see Materials and Methods section), or UV light (400 nm, see Material and Methods, and Fig. 4C) for 10 s. We averaged the stiffness obtained during cycles 1-2 (calling it stiffness “before”) and the stiffness obtained immediately after illumination, ie. during cycles 3-4 (calling it stiffness “after”) and compared these stiffness values just before and just after light excitation. We observed stiffening consistent with our previous observations, and that blue light did not induce more stiffening than indentation alone, i.e., without light excitation (21 vs 23% median increase, respectively). However, UV light led to a significantly larger 56-% stiffening. Using the same protocol, we measured a 43% increase in stiffness due to UV light in BAECs in the absence of a fluorophore (Fig. 4D).

Knowing that the first indentation led to stiffening but that letting cells rest in the dark for 10 seconds would limit the stiffening, we asked whether, by letting cells rest between each indentation, we could submit cells to several indentations in a row without stiffening them. We conducted experiments in which T cells pre-incubated with DMSO were indented twelve times in a row (“12-cycle one-color protocol” described in Figure 3E). In these experiments, after the second indentation and in-between each indentation up to 10 indentations, cells were left in the dark for 4s. As shown in Figure 4C (top, blue power 0%), the median cell stiffness was very stable over 12 indentations, increasing by only 1.2% between each cycle (Fig. 4D shows the relative stiffness over cycles 1 to 12).

Although blue light did not affect the evaluated stiffness more than indentation itself using a single exposure to 10-s blue excitation, we asked if the effect of blue light could be observed when accumulating light dose and stimulation, i.e., when subjecting each cell to a series of several consecutive 4-s excitation periods of blue light, using the 12-cycle one-color protocol (Fig. 4C, top). Using this protocol, cells stiffened more due to blue light excitation (5.9 % every cycle, Fig. 4D) than when leaving them in the dark during the 4-s periods (1.2% per cycle, Fig. 4D). Thus, although UV light has a more potent stiffening effect than blue light, blue light’s stiffening effect could be observed using several successive light excitation cycles. This showed that even without being loaded with a fluorophore, cells reacted to and stiffened when excited by blue light. Note that we used 100% power of the fluorescence excitation (blue or UV LED) in both cases, but we also checked that this corresponded to a comparable total irradiance. The blue LED was, in fact, about 3-fold more powerful (43000 mW/cm2 vs 16000 mW/cm2, Supp. Fig. 5 and Supp. Table 4) than the UV LED (using a power meter with wavelength correction), even though it led to a lower stiffening effect, highlighting the efficiency of UV light in stiffening cells.

Loading T cells with a fluorophore (Fluo-4) in the absence of excitation light did not induce more stiffening than indentation of Fluo-4-free cells in the absence of excitation light. As expected, using a 40x air objective in Fluo-4-loaded T cells in the absence of light excitation led to the same cell stiffening for repeated indentation cycles (Fig. 4C, bottom).

Overall, these results show that both the indentation and excitation of cells by blue or UV light induce and contribute to cell stiffening in our experiments, with a stronger stiffening effect in the presence of UV light than in the presence of blue light. These results also show that allowing cells to rest in the dark between indentations limits stiffening. Although these control experiments warned us of the limitations of the experiments, we will see in the following that these effects are widely overpassed by the effect of loading cells with a fluorophore and exciting this fluorophore.

### 2.5 Various cell types loaded with various fluorescent probes stiffen upon probe excitation

We first investigated the effect of different fluorophores excitation on T cell stiffness. We used our before-after protocol. As above, we call the “before” stiffness the average of the stiffness measured during cycles 1-2, and “after” stiffness the average of the stiffness measured during cycles 3-4. We loaded T cells with 2’,7’-dichlorodihydrofluorescein diacetate (DCFH-DA), Hoechst, or Fluo-4 and excited the fluorescence of these fluorophores with blue, UV, and blue light, respectively.

The DCFH-DA probe quantifies intracellular ROS. It detects ROS by oxidation of 2′,7′-dichlorodihydrofluorescein to 2′,7′-dichlorofluorescein. The diacetate form (DCFH-DA) is used because it is cell permeant. Once entered the cell, it is cleaved into non-fluorescent 2’,7’-dichlorodihydrofluorescein by esterases present in the cell. This probe is not specific to any particular ROS and is known to have an increase in fluorescence that is not linear with an increase in intracellular ROS levels [Murphy, 2022]. The Hoechst probe labels the cell nucleus through intercalation with DNA. Cell stiffness increased by approximately 10-fold due to DCFH-DA excitation, approximately 4-fold due to Hoechst excitation, and approximately 3-fold due to Fluo-4 excitation (Figure 5A). In all cases, the increase was much larger than that observed when the fluorophore was not excited (Fig. 5A). Below, Figure 6 shows an increase when exciting the CellTracker Red probe. Importantly, in all our experiments, we did not observe any noticeable changes in cell shape or contrast under bright-field illumination. Movie S3 shows an example of DCFH-DA-loaded T-cell indentation with and without fluorescence excitation.

**Figure 5.**
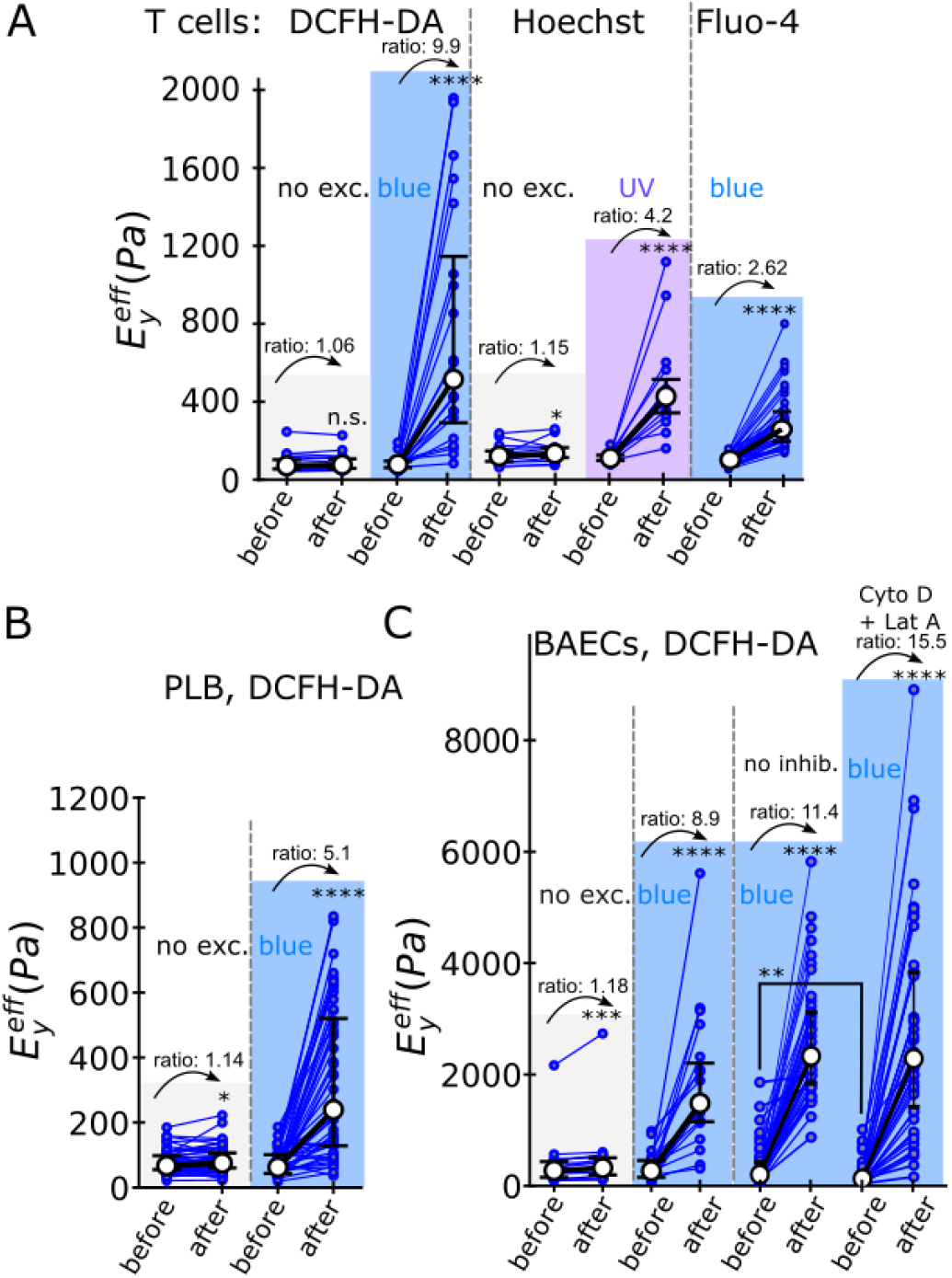
**(A)** T cells loaded with different fluorophores and submitted to the before-after protocol described in Figure 3. When loaded with DCFH-DA, Hoechst, or Fluo-4 and excited by blue, UV, or blue light, respectively, T cells become 9.9-fold, 4.2-fold, and 2.62-fold stiffer, respectively. Controls with no light excitation are shown for DCFH-DA and Hoechst loading in the absence of light excitation. They lead to 1.06- and 1.15-fold increase in stiffness, respectively. **(B)** PLB cells loaded with DCFH-DA and submitted to the before-after protocol with blue excitation become about 5-fold stiffer (median ratio 5.1). The control performed under DCFH-DA loading without light excitation is also shown. **(C)** BAECs loaded with DCFH-DA and submitted to the before-after protocol with blue excitation become about 9-fold stiffer (median ratio 8.9). The control with no light excitation and DCFH-DA loading is also shown. When incubated with Cytochalasin D and Latrunculin A inhibitors, DCFH-DA-BAECs soften, as observed in the stiffness measured before light excitation. However, upon excitation with blue light, the cell stiffness also increased by more than 11-fold.

**Figure 6.**
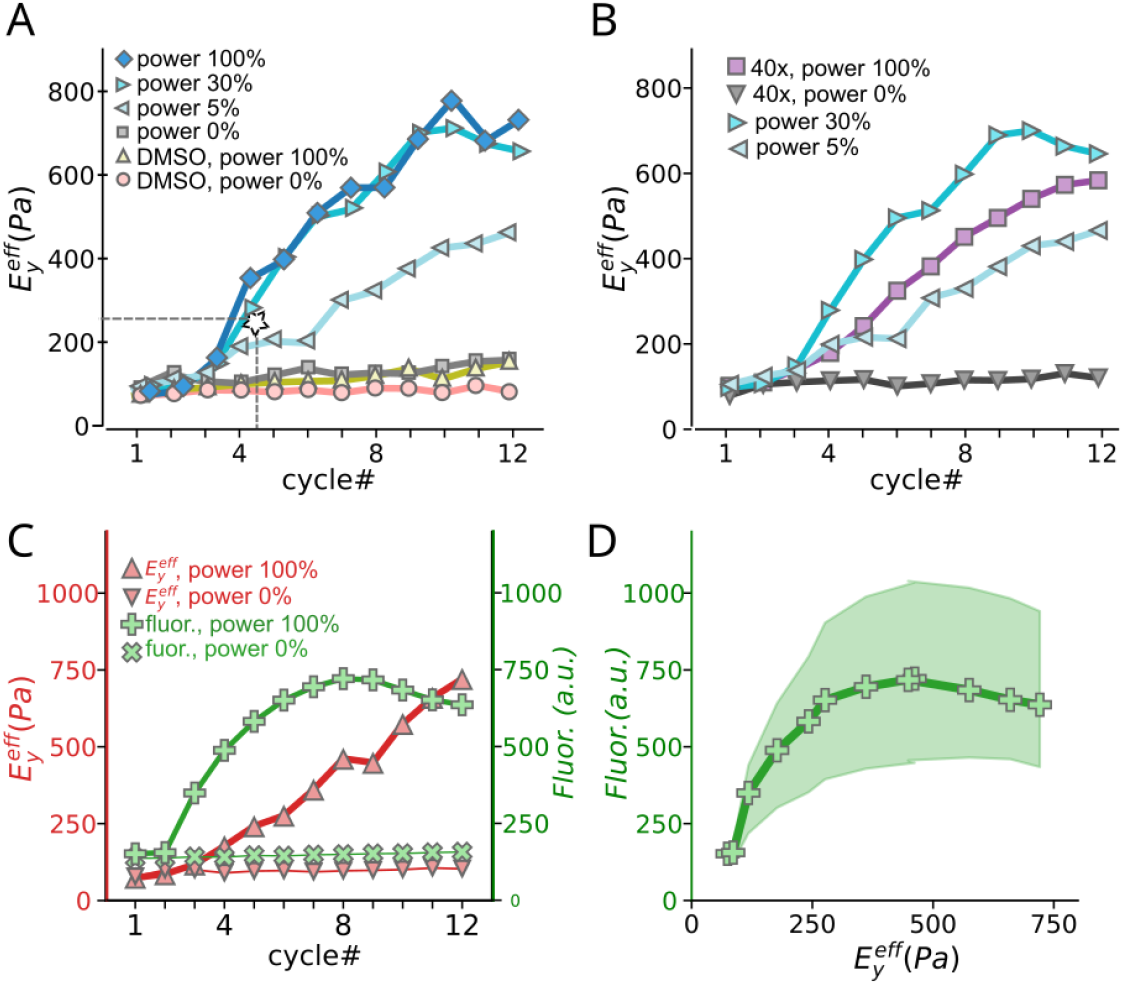
**(A)** Median stiffness vs cycle number for Fluo-4-loaded T cells in the 12-cycle one-color protocol, in which cells were indented twice without light excitation (cycles 1-2), then sequentially indented and exposed to 4 s of excitation light ten times in a row (cycles 3-12). Results obtained for different blue light powers (0%, 5%, 30%, and 100%) and control cells (incubated in DMSO in absence of Fluo-4) at 0% and 100% blue light power. For comparison, the cell stiffness values obtained under a single 10-s illumination (Figure 5A) were added (star symbol). **(B)** Median stiffness vs cycle number for Fluo-4-loaded T cells submitted to the 12-cycle one-color protocol, where a 40x air objective was used instead of the 100X oil immersion objective. For comparison, curves obtained with the 100X oil immersion objective with 5% and 30% lamp power are reproduced from panel A. **(C)** Median stiffness (red line) and fluorescence intensity (green line) vs cycle number for T cells loaded with both CellTracker Red and DCFH-DA and submitted to the 12-cycle two-color protocol. CTR was used as a phototoxic fluorophore, and DCFH-DA was used as a ROS reporter. DCFH-DA was excited with 200 ms exposure to blue light at 2% lamp power, and following this short excitation, a longer excitation (4 s) by green light at 100% power was applied to induce CTR-mediated cell stiffening. **(D)** Median fluorescence intensity *vs* median stiffness for T cells loaded with both CellTracker Red and DCFH-DA. The shaded area represents the interquartile range.

To assess the generality of this fluorescence-induced observation, we extended our investigation to other cell types: neutrophil-like PLB cells (called PLB cells herein) and (detached) bovine aortic endothelial cells (BAECs) loaded with DCFH-DA (Figure 5B-C and Movie S4). Both cell types also stiffened upon excitation of the fluorophore (PLBs: about 5-fold, BAECs: about 9-fold). We refer to the process by which cells loaded with fluorescent probes stiffen upon fluorescence excitation as photostiffening. The above results show that photostiffening is general, as it is observed in various cell types loaded with various fluorescent probes.

### 2.6 Dose accumulation in relation to the stiffening and creation of intracellular ROS

Having established that the exposure of fluorophore-loaded cells to 10 s of continuous excitation light could dramatically stiffen cells, we investigated how cell stiffness would increase upon the administration of successive doses of excitation light, as is the case when performing live movie acquisition, for example, in epifluorescence or confocal imaging. To address this question, we submitted cells to the 12-cycle one-color protocol described in Figure 3E, where cells were indented twice without light excitation (cycles 1-2), then indented and exposed to a sequence of 4 s of excitation light ten times in a row (cycles 3-12). We first studied T cells loaded with the Fluo-4 calcium probe and observed a gradual increase in cell stiffness, paralleling the increase in the accumulated dose of excitation light over time (Figure 6A). To elucidate the relationship between stiffening and intensity of light excitation, we performed a series of experiments with increasing powers of the excitation LED lamp. We used light powers of 0% (control), 5%, 30%, and 100%. In the Materials and Methods section, we present a direct measurement of the lamp power and that the output power is, as expected, reasonably linear with the requested power. The rate of stiffening observed over time increased with lamp power, but not linearly, as it saturated at 30% power, to levels similar to those when using 100% power (Fig 6A).

### 2.7 Increase in stiffness is correlated with increase in intracellular ROS concentration

The excitation of fluorescent probes is widely acknowledged for its phototoxic effects, and a key mechanistic aspect of this phenomenon is the generation of reactive oxygen species (ROS). To observe ROS production upon the exposure of cells to fluorescent light, we adapted our previously described protocol for monitoring temporal changes in cell stiffness. In our adapted protocol, CD4+ T cells were loaded with both DCFH-DA as a ROS sensor and CellTracker Red (CTR) as a phototoxic fluorophore. To mitigate potential interference due to DCFH-DA-induced cell stiffening, we employed a minimally stimulating fluorescent light dose (200 ms exposure to blue light at 2% of lamp power), which was sufficient to obtain a measurable level of fluorescence. Following this short excitation, a longer excitation (4 s) by green light was applied to induce CTR-mediated cell stiffening. As shown in Figure 6 C-D, the emission fluorescence of DCFH-DA, which serves as an indicator of intracellular ROS levels, increased together with cell stiffness, consistent with the involvement of ROS in the stiffening process, until DCFH-DA fluorescence reached a maximum and then decreased (see Movie S5 for examples). This indicates that initially, DCFH-DA reports a clear increase in ROS levels with cell stiffening, after which the DCFH-DA fluorescence level stops increasing, which might be due to bleaching or leakage.

### 2.8 DCFH-DA fluorescence dynamics

By looking at the microscope’s ocular during experiments while using the before-after protocol on T cells loaded with DCFH-DA were running, we noticed that fluorescence intensity was not constant over time, so we ran time-lapse imaging (without any mechanical measurements) on DCFH-DA-loaded T cells left at the bottom of a Petri dish or gently held by a micropipette.

In the first set of experiments, we loaded cells with DCFH-DA, gently held them in position with a micropipette, and acquired a timelapse with one image every 250 ms, exposing them for 100 ms to blue light at 100% lamp power. This resulted in a fluorescence intensity that increased over time until it peaked within seconds and then decreased again (see Movie S6 for examples of T cells). Because cells were exposed to 400 ms of blue light for every second of the time-lapse, we report the fluorescence intensity as a function of the accumulated time of exposure to blue light, as shown in Supplementary Figure 1. We observed similar behavior for BAECs in a similar setup. The accumulated time of exposure to blue light at the peak DCFH-DA fluorescence level was 5.5 s for T cells and 2.6 s for BAECs (2 donors, 25 cells, insets in Supp. Fig. 1A, C).

In a separate set of experiments, we loaded T cells with both CellTracker Red and DCFH-DA. Similar to our two-color experiments, we used CTR as the phototoxic agent and DCFH-DA at low doses to report ROS production without being too phototoxic. We thus performed 2-color time-lapse microscopy with a short exposure (50-ms) and low-intensity (10% power of the blue light) excitation of DCFH-DA and a longer (200-ms) and strong (100% power of the green light) excitation of CTR at a rate of 2 frames per second. We obtained the same qualitative behavior of DCFH-DA fluorescence level, increasing until reaching a peak and then decreasing (Supp. Fig. 1B). The cumulated green excitation time at the peak of DCFH-DA fluorescence was 16.8 s (2 donors, 21 cells, inset in Supp. Fig. 1B). This 2.7-fold longer cumulated time at the fluorescence peak, together with the fact that the 100% power green LED irradiance was about 3-fold lower than that of the 100% power blue excitation LED, indicates that reaching the peak in the DCFH-DA fluorescence level corresponded to a similar accumulated energy provided by the excitation light.

### 2.9. AFM on adherent cells

We aimed to confirm our results using a different and well-established technique in the field of cell mechanics, such as atomic force microscopy (AFM). Firm adhesion is a prerequisite for reliable AFM-based nano-indentation measurements, and we had not test photostiffening on adherent cells without detaching them. In order to complete our work on the white blood cells mechanics, and with these prerequisites in mind we chose to work with RAW macrophages [Ndao, 2020].

The excitation fluorescence intensity was measured to be much lower in the AFM setup (about 30-folds, see Supplementary Material 2). We thus partially compensated for this weaker light power by exposing cells to fluorescence for 30 s instead of 10 s in a “before-after” protocol adapted to AFM. Thus, given the weaker excitation power in this setup, we expected a small photostiffening effect. Indeed, as shown in Supplementary Figure 4, in the presence of Fluo-4, the 34% stiffening was not significantly larger than the 20-% increase due to control indentation, where Fluo-4 loaded cells were indented in the absence of blue light excitation (p-value = 0.7, Mann-Whitney U test). However, when RAW cells were loaded with DCFH-DA, excitation with blue light led to a 37% increase in stiffness that was significantly larger than the 5% increase in control experiments in which DCFH-DA-loaded cells had not been stimulated by blue light (p-value = 0.027, Mann-Whitney U test). As such, these experiments are well in line with our light-dose and dye-dependant photostiffening observations on non-adherent cells.

### 2.10 Volume vs. cortical stiffening

As explained in detail in [Markova, 2024], indenting a cell (separately) with either beads with a diameter of several microns or needles with a tip of submicron radius can help measure both the cell tension and its inner viscoelastic modulus. We asked whether photostiffening mainly affected the cell interior, but not its surface. Thus, we expected that the contact stiffness measured using a sharp tip would not increase, whereas the effective Young’s modulus measured using a large bead would do so. Thus, we performed indentation with both beads and needles on T cells loaded with Fluo-4: as shown in Supplementary Figure 5, the stiffness increased 2.6-fold after indenting with a sphere (Suppl. Fig. 5A), which was significantly more than the 1.5-fold increase in contact stiffness measured after indenting with a needle (Suppl. Fig. 5B). This suggests that the cell interior is the main contributor of the photostiffening of T cells.

### 2.11 Independence from actin cytoskeleton integrity

The actin cytoskeleton is often considered to largely determine cell stiffness [Wu, 1998; Rotsch, 2000], so we asked whether photostiffening would depend on the presence of an intact actin cytoskeleton. Thus, we destabilized the actin cytoskeleton using Cytochalasin D (CytoD) and Latrunculin A (LatA). We incubated BAECs in both cytoD (9.8µM) and LatA (1µM) and loaded these cells with DCFH-DA. As expected, inhibited cells were softer than control cells that did not incubate with inhibitors (nor with DMSO in this specific case), as shown previously (lower stiffness measured during cycle 1). However, post-illumination stiffness was unchanged (2615+/-1061 Pa vs 2831+/-1924 Pa, mean +/-SD, p = 0.97), indicating that photostiffening does not depend on the integrity of the actin cytoskeleton. As the initial stiffness decreased because of the inhibitors, the relative increase was even higher in the presence of the actin inhibitors (15.5 vs 11.4, p-value = 0.036, N =3, n=44 cells, Figure 5C).

### 2.12 Live antifade treatment does not protect cells from stiffening

DCFH-DA reported an increase in intracellular ROS during photostiffening, and to further investigate the role of ROS in this process, we tried to actively prevent the formation of ROS. Although photobleaching is not synonymous with phototoxicity, we reasoned that by limiting photobleaching using an antifade agent, we could limit phototoxicity and thus cell stiffening. We thus incubated T cells in the presence of the live antifade Prolong and loaded cells with Hoechst. The presence of Prolong did not change the stiffening of T cells upon UV illumination (Supp. Figure 2).

### 2.13 H_2_O_2_ induces T cells and BAECs stiffening

To expose cells to ROS independently of phototoxicity, we incubated T cells in the presence of 8.7 mM hydrogen peroxide (H_2_O_2_), a non-radical ROS known to diffuse into the cells across the plasma membrane. This resulted in a significant increase in cell stiffness in both T cells and detached BAEC cells (Figure 7A-B), indicating that an increased concentration of intracellular ROS alone is sufficient to induce cell stiffening.

**Figure 7.**
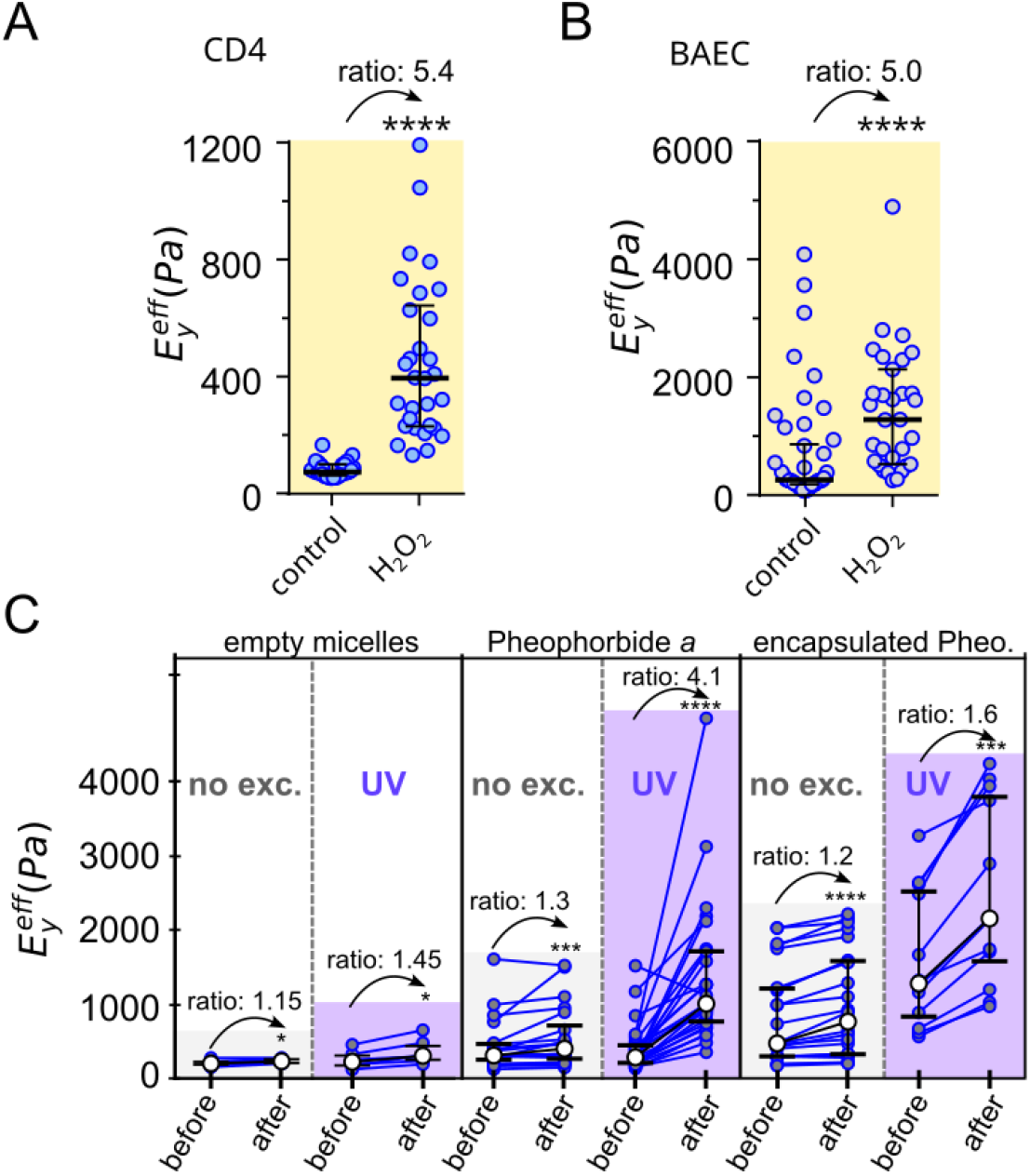
**(A)** Stiffness of T cells indented in their culture medium (control) or in culture medium with 8.7 mM hydrogen peroxide (H_2_O_2_). **(B)** Stiffness of detached BAECs cells indented in their culture medium (control) or in culture medium with 8.7 mM hydrogen peroxide (H_2_O_2_). **(C)** Detached BAECs were submitted to the before/after protocol with 10-s UV excitation, after having incubated with empty micelles (on the left: 2 independent experiments, 13 cells), Pheophorbide *a* (in the middle, 3 independent experiments, 22 cells), and encapsulated Pheophorbide *a* (on the right, 3 independent experiments, 22 cells). In all cases, the controls with no UV excitation are shown.

### 2.14 Photodynamic Therapy and Cell Stiffening

To validate our results obtained with various fluorescent probes inducing cell stiffening in multiple cell types, we opted to work with photodynamic therapy, (PDT), a clinically used therapeutic modality used for dermatological, ophthalmological, and oncological applications that relies on the local generation of reactive oxygen species (ROS) through light excitation of a photosensitizer [Verger, 2021]. PDT implies the in-situ generation of reactive oxygen species (ROS) following the illumination of a photosensitizer and transfer of its energy to the surrounding oxygen molecules [Verger, 2021]. Consequently, oxidative stress leads to cell death. Most photosensitizers tend to form aggregates in aqueous environments because of their low water solubility, which strongly decreases their ROS-photoinduced production efficiency. The main strategy to overcome this concern is to encapsulate them, especially in micelles [Demazeau, 2020]. In our experiments, when endothelial cells were subjected to incubation with the photosensitizer Pheophorbide *a* [Saide, 2020], a notable increase in cell stiffness was observed (Figure 7C). Interestingly, the photostiffening effect was much stronger when cells were incubated with encapsulated Pheophorbide *a* (see Movie S8 and Movie S9 for absence and presence of UV excitation, respectively). The encapsulation of Pheophorbide *a* in micelles enhances its cell uptake and ROS generation capabilities [Gibot2020]. Surprisingly, we also observed that empty micelles independently induced cell stiffening, even in the absence of exposure to blue light (Figure 7C). Although cell responses associated with empty polymer micelles have been reported [Brival, 2024], their stiffening effect on cells has, to our knowledge, never been reported. These results suggest that cell stiffness can serve as a useful proxy for evaluating the efficiency of photodynamic therapy, offering potential applications in monitoring therapeutic effects.

## 3. Materials and Methods

### Statistics

Experiments were performed on at least 5 cells for 3 donors or independent experiments and were pooled, unless stated otherwise. When testing if the stiffness ratio between two cycles or the stiffness before (averaging cycles 1 and 2) *vs*. after light excitation (averaging cycles 3 and 4) was larger than one, no normal distribution of the ratio was assumed, and a Mann-Whitney test was used. When testing whether stiffness increased between two cycles or when comparing “before” and “after” values, no normal distribution was assumed, and a Wilcoxon matched-pairs signed-rank test was used. Labels for *p*-values are: * for p<0.05, ** for p<0.01, *** for p<0.001, and **** for p <0.0001.

### Cells

#### T cells

The purification details of human CD4+ T cells can be found elsewhere [Zak, 2021]. Experiments were performed at room temperature in RPMI 1640 medium without phenol (GIBCO ref. 11835-030 500 mL) supplemented with L Glutamine, 10% heat-inactivated fetal bovine serum (FBS), and 1% penicillin-streptomycin (all purchased from Thermo Fisher Scientific, Waltham, MA) and filtered with 0.22-µm diameter pores (Merck Millipore, Burlington, MA).

#### PLB cells

The human acute myeloid leukemia cell line PLB-985 (called PLB cells herein) is a subline of HL-60 cells. PLB cells were cultured in RPMI 1640 medium with GlutaMax, 10% heat-inactivated FBS, and 1% penicillin-streptomycin. Cells were passaged twice a week and differentiated into a neutrophil-like phenotype by adding 1.25% (v/v) dimethyl sulfoxide (Sigma-Aldrich, St. Louis, MO) to the cell suspension on the first day after passage and a second time 3 days after changing the culture medium. Differentiated PLB-985 cells remain in suspension and exhibit neutrophil-like properties. Differentiated PLB cells were used after 5 or 6 days of differentiation. As for T cells, experiments were performed at room temperature in complete RPMI 1640 medium without phenol red.

#### Endothelial cells

Bovine aortic endothelial cells (BAECs) were used between passages 4 and 7. As previously described [Guillou, 2016; Zak, 2021], BAECs were cultured at 37 °C and 5% CO2 in Dulbecco’s Modified Eagle’s Medium (DMEM, Thermo Fisher Scientific) supplemented with 10% heat-inactivated fœtal bovine serum and 1% penicillin/streptomycin. The cells were passed two to three times a week. Before the experiments, a flask of BAECs was trypsinized with trypLE (Thermo Fisher Scientific). Cells were resuspended in complete DMEM and left for 30 min on a rotating wheel in aluminum foil before injection into the bottom of a glass-bottom Petri dish (Fluorodish; World Precision Instruments, Hitchin, UK) in 3 mL of DMEM/F-12 medium without phenol red (Fisher Scientific, ref. 11039021) supplemented with 10% heat-inactivated fœtal bovine serum and 1% penicillin/streptomycin, and filtered with 0.22-µm diameter pores.

#### RAW cells

RAW 264.7 macrophage cells were obtained from the European Collection of Cell Cultures (ECACC, Salisbury, UK). The cells were cultured at 37°C with 5% CO2 in RPMI 1640 medium (Life Technologies, Aubin, France) supplemented with 10% heat-inactivated fetal bovine serum (FBS, Life Technologies) and 2 mM L-glutamine. Cells were monitored regularly and subcultured when they reached approximately 80% confluence, typically every 2-3 days. For subculturing, cells were gently detached using a cell scraper, centrifuged at 300 RPM for 3 min, and reseeded at a 1:10 dilution in fresh culture medium. Before the AFM experiments, cells were detached using classical protocols [Ndao, 2020], seeded in glass-bottom Petri dishes (Fluorodish) to a low density, and allowed to adhere overnight. The experiments were performed on the following day, with the cells well separated on the surface.

### Reagents

2’,7’-Dichlorofluorescein diacetate (Sigma) was aliquoted at 30 mM in DMSO; final concentration during the 20-min incubation was 10 µM for both microindentation and AFM experiments. Fluo-4, AM (ThermoFisher) was aliquoted at 1 mM in DMSO; final concentration during the 20-min incubation was 2 µM for microindentation and 6 µM for AFM experiments. CellTracker Red CMTPX (Invitrogen, ThermoFisher, ref. C34552) was used at a final concentration of 2mM during a 20-min incubation. In the text, It is called CellTracker Red or CTR. Hoechst 33342 (invitrogen, ThermoFisher) was used with a dilution of 1:2000 of the stock solution (10mg/ml) during the 20-min incubation. Cytochalasin D (Sigma, ref. C2618-200µL Ready Made Solution, from Zygosporium mansonii, 5 mg/mL in DMSO, 0.2 μm filtered) was used at a final concentration of 9.8 µM during incubation and experiments. Latrunculin A (Sigma ref. 428026-50UG) was used at final concentration 1 µM during incubation and experiments. H_2_O_2_ was purchased from sigma (ref. 216763-100mL, 30% H_2_O_2_ in H_2_O) and used at a final concentration 8.7 mM in complete DMEM/F12 (with 10% serum) or complete RPMI (with 10% serum) for BAECs and T cells, respectively. T cells were incubated with 100-fold diluted ProLong Live Antifade Reagent (Thermofisher) stock solution for 2 h in an incubator at 37°C and 5% CO_2_. BAECs were incubated in the dark with empty poly(ethylene oxide)-block-poly(ε-caprolactone) PEO_5000_-PCL_4000_ micelles (Polymer Source), free Pheophorbide *a* (Frontier Scientific), or Pheo-loaded PEO-PCL micelles for 30 min at a Pheo concentration of 1.65 μM [Gibot, 2020]. For DMSO control experiments, cells were incubated in 0.2% DMSO.

### Profile microindentation

#### Principle

We have previously described in detail the profile microindentation technique [Husson, 2023], which allows sideways observation of cells under the microscope during indentation. It consists of holding a cell with a stiff micropipette solidary to a piezo single-axis translation stage. The cell is then translated against a flexible micropipette with known bending stiffness. The latter, termed microindenter, has a tip with a spherical or needle shape. By detecting the microindenter tip’s position on a live camera image, the deflection of the microindenter can be measured, from which the force applied to the cell can be deduced. To apply the desired force, the position of the piezo stage is adjusted via a live feedback loop. From the difference between the piezo position and the microindenter’s tip, the indentation of the cell is calculated.

#### Micropipette and microneedle preparation

Stiff micropipettes were prepared as described previously [Guillou 2016; Sawicka, 2017; Husson, 2023] from borosilicate glass capillaries (Harvard Apparatus, Holliston, MA, USA) pulled with a micropipette puller (P-97, Sutter Instruments, Novato, CA, USA) and cut with a microforge (MF-200, World Precision Instruments) to obtain the desired tip diameter (3.5 µm for T cells and PLB cells, 6-9 µm for detached ECs). The Micropipettes were bent at 45° with a second microforge (MF-900, Narishige, Tokyo, Japan). The extremities of flexible micropipettes were melted into a glass bead or a microneedle (Fig. 7B). The radii of both spherical indenters and microneedles were measured optically with an accuracy of typically 3 pixels (about 0.2 µm). As previously described [Sawicka, 2017; Husson, 2023], the bending stiffness of flexible micropipettes was calibrated by pressing them against standard microindenters of known stiffness. The stiffness of the standard microindenters was calibrated using a commercial force probe (model 406A with a force range of 0–500 nN; Aurora Scientific, Aurora, ON, Canada). Microindenters and microneedles had a typical bending stiffness of 0.6 nN/µm when working with T cells and PLB cells, and 1-2 nN/µm when working with BAECs. Microindenters with a needle-like tip had a typical stiffness of 0.1 nN/µm.

#### Setup

As described previously [Husson, 2023], an inverted microscope (TiE, Nikon Instruments, Tokyo, Japan) was placed on an air suspension table (Newport, Irvine, CA, USA) equipped with a Flash 4.0 CMOS camera (Hamamatsu Photonics, Hamamatsu City, Japan), a 100x oil immersion 1.3 numerical aperture (NA) objective (Nikon Instruments), and a 40x air objective (Nikon Instruments). The stiff micropipette is attached to a piezoelectric single-axis translation stage (Physik Instrumente, Karlsruhe, Germany), which is itself placed on a custom-made adapter piece attached to a motorized micromanipulator (MP-285; Sutter Instruments) for 3D placement of the cell before any microindentation. The microindenter is attached to a non-motorized micropositioner (Thorlabs, Newton, NJ). The micropipette and microindenter are then plunged into a glass-bottom Petri dish (Fluorodish, World Precision Instruments). The stiff micropipette is connected to a water-filled reservoir (made of a 50-mL syringe body) whose height is adjusted to control the hydrostatic pressure inside the micropipette [Husson, 2023]. A typical aspiration of a 5-mm H_2_O difference (an aspiration pressure of about 50 Pa) was set to gently hold the cells. Experiments were performed at room temperature to avoid thermal drift. LEDs were used as the fluorescence light sources (CoolLED pE-300 ultra, Micromecanique, Saint-Germain-en-Laye, France). When working with both DCFH-DA and CTR, a GFP-DSRED fluorescence filter block was used; when working with DCFH-DA alone and Fluo-4, a FITC fluorescence block was used; and when working with Hoechst, a DAPI fluorescence block was used. The wavelengths of the LED light source and their respective irradiance are given below.

#### Live detection

The experiment is automated by a Matlab (Mathworks, Natick, MA) code that controls the piezo stage position and the Micromanager software [Edelstein, 2014] that controls the microscope camera. The position of the microindenter tip is detected by analyzing a line crossing the shaft of the microindenter on the live-microscopy image acquired by the camera (Figure 3A). The intensity profile along this line is analyzed at an acquisition frequency of about 400 Hz and cross-correlated to a reference intensity profile recorded at the beginning of an experiment. The position of the microindenter is measured with a subpixel accuracy of 20-30 nm (one pixel represents 65 nm using the 100x oil objective) [Markova, 2024]. Images of the region of interest showing the micropipettes, the cell, and the bead at a typical frequency of 1 frame per second. Some supplementary movies were acquired at 10 frames per second.

### Measurement of the protrusion emitted by T cells during activation

The setup was described previously [Sawicka, 2017; Zak, 2021; Zucchetti, 2021]. In brief, the same system as that used for profile microindentation was used, except that the microindenter was replaced by a flexible micropipette with a bending stiffness of 0.2-0.5 nN/µm that aspirated a 4.5-µm activating microbead (Dynabeads Human T-Activator CD3/CD28 for T Cell Expansion and Activation, Gibco, Thermo Fisher Scientific, ref. 11131D). Once a preset compressive force of 200 pN was reached, the force applied to the cells was set to 0, and the protrusion emitted by T cells was free to expand. Its length was measured as a function of time. See Movie S2.

### Deformation of T cells against a rigid wall

The rigid wall was made of a solid glass rod that was cut using a microforge (MF-900, Narishige, Tokyo, Japan) to form a disk of about 20 µm in diameter. An additional 45° bent was added using the microforge. See Movie S1.

### Atomic force microscopy of RAW macrophages

An AFM head (Nanowizard 4XP, JPK Instruments/Bruker) was mounted on an inverted microscope (Zeiss Axiovert 200) equipped with a 40x NA0.9 objective, a fluorescence excitation diode (CoolLed pE-300), a CoolSnap HQ2 camera (Photometrics), and placed on an active damping table (Halcyonics). The measurements were conducted in closed-loop and constant height-feedback modes, and all experiments were performed at room temperature (to fit the micropipette experiments). We used 5µm sphere tip levers (MLCT-SPH-5UM, Bruker, nominal spring constant 10 pN/nm). The sensitivity of the optical lever system was calibrated on a glass substrate, and the spring constant was measured using the built-in thermal noise method before each experiment at room temperature. Fluorescence images were captured using a FITC cube, and bright-field images were used to select regions of interest. Image acquisition was performed using Micromanager software (version 1.4.22). In the AFM experiments, the lower excitation light power was partially compensated by illuminating cells for 30 s instead of 10 s in the microindentation experiments. More details on the measurement protocol and data analysis are given in Supplementary Material 3.

### Power meter

In both the profile microindentation and AFM setups, the power at the specimen was measured using a microscope slide power sensor (S170C, Thorlabs), connected to a digital optical power meter console (PM100D, Thorlabs), including correction for excitation wavelength. The exposed surface area was controlled using the microscope diaphragm. See Supplementary Material 1 for the measured power depending on the wavelength and lamp power in the profile microindentation setup, and Supplementary Material 2 for the measured power depending on the lamp power in the AFM setup.

## 4. Discussion

In this study, we demonstrated that upon excitation for several seconds with an LED light source, fluorophore-loaded cells become much stiffer within seconds. This photostiffening effect explains why T cells loaded with the Fluo-4 calcium probe stop emitting a protrusion within seconds when excitation light is turned on or modify, even stop, their migration or force exertion on surfaces. We observed photostiffening in different cell types (T cells, endothelial cells, neutrophils and macrophages) and with distinct fluorophores (Fluo-4, Celltracker Red CMTPX, Hoechst, and DCFH-DA). Furthermore, the cell stiffness increased with increasing exposure time to excitation light and increasing power of the light source for a given excitation duration. Our profile microindentation showed that cell indentation alone led to moderate cell stiffening. In the absence of a fluorophore, excitation with blue or UV light led to moderate cell stiffening, which was much less than that observed when a fluorophore was excited. Furthermore, we used both sharp and blunt indenters to demonstrate that stiffening occurred not only at the cell cortex level but also in the cell interior. We also correlated the increase in cell stiffness with the production of intracellular ROS and reproduced cell stiffening by incubating cells in H_2_O_2_. Excitation of the photosensitizer Pheophorbide *a*, which induces specific types of ROS (singlet oxygen species), also led to cell stiffening.

Phototoxicity has been described for several decades [Wang, 1976], including the toxic effects of blue and UV light in cells in the absence of a fluorophore and in cells expressing a fluorescent protein in which an increased production of ROS upon fluorescence excitation was observed [Cheng, 2016; Diaspro, 2006; König, 1996; Laissue, 2017; Roehlecke, 2009; Wäldchen, 2015]. The culture medium can itself be a source of phototoxic molecules generated by light excitation [Stockley, 2017]. There is abundant literature on phototoxicity caused by a wide range of nature and quantities of ROS linked to the variety of fluorophores employed and to the varying irradiance of the sample used by different fluorescence imaging modalities or time sequences (epifluorescence, laser scanning- or spinning disc-confocal microscopy, light sheet and super-resolution microscopy) [Cole, 2014]. Furthermore, although the light dose is relevant for setting the photodamage level, the delivery method of the dose is also important. Natural ROS scavenger molecules are present in cells and can cope with ROS generation up to a certain rate of generation [Dixit and Cyr, 2003; Tinevez, 2012], a rate that we probably overpassed even in our experiments using the Prolong antifade agent.

Despite the existing literature, studies on the effects of phototoxicity on cell stiffness are scarce and rarely systematic. In this study, we systematically quantified the photostiffening resulting from several types of fluorophores and cell types. Some studies used ROS-inducing molecules, independent of stimulation by light, and observed an increase in stiffness (up to 2.6- and 100-fold for oridonin [Pi, 2015] and other chemotherapies [Lam, 2007], respectively). In apparent contradiction, some studies reported softening in cells exposed to ROS-inducing carbon-based nanomaterials [Pastrana, 2019], UVB radiation [Sobiepanek, 2016], and UVC radiation in the presence of nanocrystalline titanium dioxide [Vileno, 2007]. In these studies, leading to cell softening, the dose delivered to cells was relatively small compared with ours (18-100 mJ/cm^2^), and in some cases, cells were left for 24 h for recovery. We speculate that this induces a relatively small amount of ROS, which might be able to dismantle the cytoskeleton while creating a limited amount of covalent cross links that cannot compensate for the loss in structure induced, thus explaining the decrease in stiffness. In this study, we showed in T cells that the cell membrane proximal structures are less affected than those of the cell interior by photostiffening. In studies probing cell stiffness using sharp AFM tips, we expect that only th e cell surface is probed [Markova, 2024], which might soften under a limited light dose. It is thus possible that some studies have observed softening only at the cell surface, which might reflect actomyosin cortex destabilization, whereas the cell interior/cytosol or other cytoskeletal component might exhibit stiffening, possibly through the creation of covalent bonds between proteins. Note that although we only considered an effective Young’s modulus in the present study, others have reported an increase in intracellular viscosity using porphyrin-dimer-based molecular rotors [Kuimova, 2009] or photosensitizing chlorins [Aubertin, 2013]. It has been shown in many cell types, including leukocytes [Zak, 2021], that parallel increases in cell stiffness and viscosity can be expected in cells, thus supporting the idea that the intracellular medium might stiffen, even though the cell surface might not.

Our observation of light-induced “freezing” of Fluo-4-loaded T cells during activation echoes a study in which part of illuminated cells with and without fluorophores became so-called “frozen cells” i.e. completely immobile cells attached to the surface [Wäldchen, 2015]. However, Wäldchen *et al*. used a massive dose of fluorescence compared with ours, several 100 kJ/cm^2^, and although the presence of a fluorophore made cells more sensitive to irradiation, it only shifted their I50 irradiation Intensity (value of light intensity where 50% of the irradiated cells died after imaging at 514 nm for 240 s) by 20-25%. This is quite in contrast with our T cell activation experiments, where only 2-s excitation was enough to stop cell protrusions, i.e., about 80 J/cm^2^, three orders of magnitude lower, while in controlled cells loaded with Fluo-4 but not excited with blue light, cells proceeded emitting their characteristic protrusion unperturbed. In our experiments on T cells and BAECs loaded with DCFH-DA, about 400 J/cm^2^ led to severe stiffening: thus, at doses still orders of magnitude lower than those delivered by Wäldchen et al. This highlights the potential variability depending on the fluorophore, cell type (adherent or not, cell line or primary cells), and its capacity to repair or compensate for photodamage. Along this line, in the same study, Wäldchen et al. showed that different cell lines show different resistance to photodamage, with HeLa being remarkably more resistant than U2OS and COS-7 lines. They also explored the dependence of the excitation wavelength (from 405 to 639 nm), highlighting the damaging efficiency at lower wavelengths. Note that most in vitro experiments involve illuminating cells that are in the relative dark in the body. If one considers the solar irradiance of about 100 mW/cm^2^ (central Europe [Stelzer, 2014]), 10 min illumination corresponds to 60 J/cm^2^, and even though part of the sun’s irradiation spectrum falls within infrared, this dose is comparable to the maximal dose at 405 nm before phototoxicity reported by Wäldchen et al. [Wäldchen, 2015]. Fluorophore-loaded cells left for tens of minutes in a Petri dish and continuously exposed to ambient light might thus be affected by this unwanted illumination, aside from the expected dye fading. As a comparison, one can also consider the typical light dose of about 100 J/cm^2^, which seems enough to induce severe mechanical effects, which is obtained using our setup for 10-s illumination at 100% lamp power. This total dose can be delivered using 2500 snapshots of 200-ms exposure at 2% lamp power, corresponding to a timelapse of only 42 minutes for snapshots taken every second. Although this estimate discards the buffering capacity of the cells mentioned above, it highlights the caution to be taken when performing live-cell fluorescence experiments for imaging dynamical and/or long-term cell behaviors.

Using our 12-cycle protocol, we observed increased stiffness with ROS production. As expected, the rate of stiffening with accumulated dose increased at higher lamp power until reaching a plateau, as the result obtained at 30% power was the same as that obtained at 100% power. This suggests that although the relationship is unlikely to be linear, stiffness can quantitatively indicate the level of photodamage. The relationship between cell stiffness and phototoxicity opens an interesting avenue for assessing the efficiency of photodynamic therapy, which uses phototoxicity on purpose. Measuring mechanical changes in cells could be a way to rapidly assess the efficiency of different photosensitizers depending on their formulation, when tested on both healthy and patient-derived cells. Cell stiffening may also occur in other cancer therapies. Radiotherapy induces ROS production and subsequent ROS-induced cancer cell damage. The emerging FLASH radiotherapy delivers the same dose of radiation in much shorter time and appears to be less damaging to the tissue surrounding the cancer by unresolved mechanisms [Tang, 2024]. Our data suggest that cell stiffening may be a side effect of radiotherapy with yet unknown consequences. Changes in cell stiffness could also be used in this context to assess the efficacy of the treatment or the absence of adverse effects in nearby tissue.

In this study, we also artificially induced ROS generation via fluorophore excitation; however, it is unclear whether cells can use ROS to modulate their mechanical properties in physiological contexts. It has been shown in plants that reactive oxygen species modulate the tip growth of root hairs by affecting the wall properties [Monshausen, 2007]. However, to our knowledge, whether ROS can regulate cell mechanical properties under physiological conditions in mammalian cells is yet to be clarified.

In conclusion, aside from reminding that it is crucial to perform controls when using fluorescence and that—as previously stated by others—a trade-off between sample damage and the signal-to-noise ratio must always be determined [Laissue 2017], this study shows that photostiffening could be exploited as a new method for rapidly quantifying the phototoxicity of a given dye under defined illumination conditions.

## Acknowledgements

The authors thank C. Hivroz for providing T cells, A. Babataheri and A. Castagnino for expansion of Endothelial cells. JH thanks C. Stringari, B. Lordon, G. Gallot, R. Lasserre, and S. Drevensek for fruitful discussion. This work has received financial support from CNRS through the MITI interdisciplinary programs, from École Polytechnique, and from the AXA Research Fund. J. EH. is supported by a PhD funding stemming from the ANR “BreakInTheWall” (ANR-22-CE35-0008). JH and PHP acknowledge the funding from ANR “Criticality” (ANR-23-CE30-0006). LG acknowledges the funding from ANR “TeraCellATR” (ANR-21-CE42-0018). J. EH. and PHP acknowledge the technical support from the shared cell culture platform (PCC) and practical help from Martine Pelicot-Biarnes.

## Author contributions

P.-H.P. and J.H. designed experiments. E.G., J.EH., F.B.M., and J.H. performed experiments. E.G., J.EH., F.B.M., P.-H.P., and J.H. analyzed the data. T.C., H.J., S.D., P.-H.P., O.N., and L.G. contributed material. E.G., J.EH. and J.H. wrote the manuscript with helpful critical reading by the other authors. For the purpose of open access, the author has applied a Creative Commons Attribution (CC BY) license to any Author Accepted Manuscript version arising from this submission.

## References

Ahmad, S., Khan, H., Shahab, U., Rehman, S., Rafi, Z., Khan, M. Y., Ansari, A., Siddiqui, Z., Ashraf, J. M., Abdullah, S. M. S., et al. (2017). Protein oxidation: An overview of metabolism of sulphur containing amino acid, cysteine. Frontiers in Bioscience (Scholarly Edition), 9, 71–87. 10.2741/s474

Alghamdi, R. A., Exposito-Rodriguez, M., Mullineaux, P. M., Brooke, G. N., & Laissue, P. P. (2021). Assessing phototoxicity in a mammalian cell line: How low levels of blue light affect motility in PC3 cells. Frontiers in Cell and Developmental Biology, 9, 738786. 10.3389/fcell.2021.738786

Aubertin, K., Bonneau, S., Silva, A. K. A., Bacri, J. C., Gallet, F., & Wilhelm, C. (2013). Impact of photosensitizers activation on intracellular trafficking and viscosity. PLoS ONE, 8(12), e84850. 10.1371/journal.pone.0084850

Brival R, Ghafari N, Mingotaud AF, Fourquaux I, Gilard V, Collin F, Vicendo P, Balayssac S, Gibot L. (2024). Encapsulation of photosensitizer worsen cell responses after photodynamic therapy protocol and polymer micelles act as biomodulators on their own. Int J Pharm; 663:124589. doi: 10.1016/j.ijpharm.2024.124589.

Cheng, K. P., Kiernan, E. A., Eliceiri, K. W., Williams, J. C., & Watters, J. J. (2016). Blue light modulates murine microglial gene expression in the absence of optogenetic protein expression. Scientific Reports, 6, 21172.

Cole R. Live-cell imaging. Cell Adh Migr. 2014;8(5):452–9. doi: 10.4161/cam.28348.

Demazeau M, Gibot L, Mingotaud AF, Vicendo P, Roux C, Lonetti B. Rational design of block copolymer self-assemblies in photodynamic therapy. Beilstein J Nanotechnol. 2020 Jan 15; 11:180–212. doi: 10.3762/bjnano.11.15.

Diaspro, A., Chirico, G., Usai, C., Ramoino P., and Dobrucki J. (2006). Photobleaching. In “Handbook of Biological Confocal Microscopy,” (J. B. Pawley, ed.) 3rd edn, pp. 690–702. doi: 10.1007/978-0-387-45524-2_39

Dixit, R., and Cyr, R., 2003, Cell damage and reactive oxygen species production induced by fluorescence microscopy: Effect on mitosis and guidelines for non-invasive fluorescence microscopy, Plant J. 36: 280–290.

Douthwright, S., and Sluder, G. (2017). Live Cell Imaging: Assessing the Phototoxicity of 488 and 546 nm Light and Methods to Alleviate it. J Cell Physiol 232, 2461–2468. 10.1002/jcp.25588.

Edelstein AD, Tsuchida MA, Amodaj N, Pinkard H, Vale RD, Stuurman N. Advanced methods of microscope control using μManager software. J Biol Methods. 2014;1(2):e10. doi: 10.14440/jbm.2014.36. PMID: 25606571; PMCID: PMC4297649.

Feng, Y., Wang, J., Hu, H., and Yang, C. (2022). Effect of oxidative modification by reactive oxygen species (ROS) on the aggregation of whey protein concentrate (WPC). Food Hydrocolloids 123, 107189. 10.1016/j.foodhyd.2021.107189.

Galie, P.A., Georges, P.C., and Janmey, P.A. (2022). How do cells stiffen? Biochem J 479, 1825–1842. 10.1042/BCJ20210806.

Gibot, L., Demazeau, M., Pimienta, V., Mingotaud, A.-F., Vicendo, P., Collin, F., Martins-Froment, N., Dejean, S., Nottelet, B., Roux, C., et al. (2020). Role of Polymer Micelles in the Delivery of Photodynamic Therapy Agent to Liposomes and Cells. Cancers (Basel) 12, 384. 10.3390/cancers12020384.

Guillou L, Babataheri A, Puech PH, Barakat AI, Husson J. Dynamic monitoring of cell mechanical properties using profile microindentation. Sci Rep. 2016 Feb 9; 6:21529. doi: 10.1038/srep21529.

Husson, J. (2023). Measuring Cell Mechanical Properties Using Microindentation. Methods Mol Biol 2600, 3–23. 10.1007/978-1-0716-2851-5_1.

Icha, J., Weber, M., Waters, J.C., and Norden, C. (2017). Phototoxicity in live fluorescence microscopy, and how to avoid it. Bioessays 39. 10.1002/bies.201700003.

König K, Tadir Y, Patrizio P, Berns MW, Tromberg BJ. Effects of ultraviolet exposure and near infrared laser tweezers on human spermatozoa. Hum Reprod. 1996 Oct;11(10):2162–4. doi: 10.1093/oxfordjournals.humrep.a019069. PMID: 8943522.

Kuimova, M.K., Botchway, S.W., Parker, A.W., Balaz, M., Collins, H.A., Anderson, H.L., Suhling, K., and Ogilby, P.R. (2009). Imaging intracellular viscosity of a single cell during photoinduced cell death. Nature Chem 1, 69–73. 10.1038/nchem.120.

Laissue, P., Alghamdi, R., Tomancak, P. et al. Assessing phototoxicity in live fluorescence imaging. Nat Methods 14, 657–661 (2017). 10.1038/nmeth.4344.

Lam, W.A., Rosenbluth, M.J., and Fletcher, D.A. (2007). Chemotherapy exposure increases leukemia cell stiffness. Blood 109, 3505–3508. 10.1182/blood-2006-08-043570.

Markova O, Clanet C, Husson J. Quantifying both viscoelasticity and surface tension: Why sharp tips overestimate cell stiffness. Biophys J. 2024 Jan 16;123(2):210–220. doi: 10.1016/j.bpj.2023.12.008.

Milzani, A., DalleDonne, I., and Colombo, R. (1997). Prolonged oxidative stress on actin. Arch Biochem Biophys 339, 267–274. 10.1006/abbi.1996.9847.

Monshausen GB, Bibikova TN, Messerli MA, Shi C, Gilroy S. Oscillations in extracellular pH and reactive oxygen species modulate tip growth of Arabidopsis root hairs. Proc Natl Acad Sci U S A. 2007 Dec 26;104(52):20996–1001. doi: 10.1073/pnas.0708586104. Epub 2007 Dec 13. PMID: 18079291; PMCID: PMC2409255.

Mubaid, F., and Brown, C.M. (2017). Less is More: Longer Exposure Times with Low Light Intensity is Less Photo-Toxic. Microscopy Today 25, 26–35. 10.1017/S1551929517000980.

Murphy, M.P., Bayir, H., Belousov, V., Chang, C.J., Davies, K.J.A., Davies, M.J., Dick, T.P., Finkel, T., Forman, H.J., Janssen-Heininger, Y., et al. (2022). Guidelines for measuring reactive oxygen species and oxidative damage in cells and in vivo. Nat Metab 4, 651–662. 10.1038/s42255-022-00591-z.

Ndao O, Puech PH, Bérard C, Limozin L, Rabhi S, Azas N, Dubey JP, Dumètre A. Dynamics of Toxoplasma gondii Oocyst Phagocytosis by Macrophages. Front Cell Infect Microbiol. 2020 May 19; 10:207. doi: 10.3389/fcimb.2020.00207.

Olinski, R., Nackerdien, Z., and Dizdaroglu, M. (1992). DNA-protein cross-linking between thymine and tyrosine in chromatin of gamma-irradiated or H2O2-treated cultured human cells. Arch Biochem Biophys 297, 139–143. 10.1016/0003-9861(92)90651-c.

Pastrana HF, Cartagena-Rivera AX, Raman A, Ávila A. Evaluation of the elastic Young’s modulus and cytotoxicity variations in fibroblasts exposed to carbon-based nanomaterials. J Nanobiotechnology. 2019 Feb 23;17(1):32. doi: 10.1186/s12951-019-0460-8. PMID: 30797235; PMCID: PMC6387485.

Pi J, Cai H, Jin H, Yang F, Jiang J, Wu A, et al. (2015) Qualitative and Quantitative Analysis of ROS-Mediated Oridonin-Induced Oesophageal Cancer KYSE-150 Cell Apoptosis by Atomic Force Microscopy. PLoS ONE 10(10): e0140935. doi:10.1371/journal.pone.0140935.

Roehlecke C, Schaller A, Knels L, Funk RH. The influence of sublethal blue light exposure on human RPE cells. Mol Vis. 2009 Sep 21; 15:1929–38. PMID: 19784391; PMCID: PMC2751800.

Rotsch C, Radmacher M. Drug-induced changes of cytoskeletal structure and mechanics in fibroblasts: an atomic force microscopy study. Biophys J. 2000 Jan;78(1):520–35. doi: 10.1016/S0006-3495(00)76614-8.

Sawicka A, Babataheri A, Dogniaux S, Barakat AI, Gonzalez-Rodriguez D, Hivroz C, Husson J. Micropipette force probe to quantify single-cell force generation: application to T-cell activation. Mol Biol Cell. 2017 Nov 7;28(23):3229–3239. doi: 10.1091/mbc.E17-06-0385.

Saide A, Lauritano C, Ianora A. Pheophorbide a: State of the Art. Mar Drugs. 2020 May 14;18(5):257. doi: 10.3390/md18050257.

Sies, H., Belousov, V.V., Chandel, N.S. et al. Defining roles of specific reactive oxygen species (ROS) in cell biology and physiology. Nat Rev Mol Cell Biol 23, 499–515 (2022). 10.1038/s41580-022-00456-z

Sindhu, A., Janick-Buckner, D., Buckner, B., Gray, J., Zehr, U., Dilkes, B.P., and Johal, G.S. (2018). Propagation of cell death in dropdead1, a sorghum ortholog of the maize lls1 mutant. PLoS One 13, e0201359. 10.1371/journal.pone.0201359.

Sobiepanek A, Milner-Krawczyk M, Bobecka-Wesołowska K, Kobiela T. The effect of delphinidin on the mechanical properties of keratinocytes exposed to UVB radiation. J Photochem Photobiol B. 2016 Nov; 164:264–270. doi: 10.1016/j.jphotobiol.2016.09.038.

Stelzer EH. Light-sheet fluorescence microscopy for quantitative biology. Nat Methods. 2015 Jan;12(1):23–6. doi: 10.1038/nmeth.3219. PMID: 25549266.

Stockley JH, Evans K, Matthey M, Volbracht K, Agathou S, Mukanowa J, Burrone J, Káradóttir RT. Surpassing light-induced cell damage in vitro with novel cell culture media. Sci Rep. 2017 Apr 12;7(1):849. doi: 10.1038/s41598-017-00829-x. PMID: 28405003; PMCID: PMC5429800.

Tang R, Yin J, Liu Y, Xue J. FLASH radiotherapy: A new milestone in the field of cancer radiotherapy. Cancer Lett. 2024 Apr 10; 587:216651. doi: 10.1016/j.canlet.2024.216651.

Tinevez JY, Dragavon J, Baba-Aissa L, Roux P, Perret E, Canivet A, Galy V, Shorte S. A quantitative method for measuring phototoxicity of a live cell imaging microscope. Methods Enzymol. 2012; 506:291–309. doi: 10.1016/B978-0-12-391856-7.00039-1. PMID: 22341230.

Verger A, Brandhonneur N, Molard Y, Cordier S, Kowouvi K, Amela-Cortes M, Dollo G. From molecules to nanovectors: Current state of the art and applications of photosensitizers in photodynamic therapy. Int J Pharm. 2021 Jul 15; 604:120763. doi: 10.1016/j.ijpharm.2021.120763.

Vileno, B., Lekka, M., Sienkiewicz, A., Jeney, S., Stoessel, G., Lekki, J., Forró, L., and Stachura, Z. (2007). Stiffness Alterations of Single Cells Induced by UV in the Presence of NanoTiO 2. Environ. Sci. Technol. 41, 5149–5153. 10.1021/es0629561.

Wäldchen S, Lehmann J, Klein T, van de Linde S, Sauer M. Light-induced cell damage in live-cell super-resolution microscopy. Sci Rep. 2015 Oct 20; 5:15348. doi: 10.1038/srep15348.

Wang RJ. Effect of room fluorescent light on the deterioration of tissue culture medium. In Vitro. 1976 Jan;12(1):19–22. doi: 10.1007/BF02832788.

Wu, H.W., Kuhn, T. and Moy, V.T. (1998), Mechanical properties of L929 cells measured by atomic force microscopy: Effects of anticytoskeletal drugs and membrane crosslinking. Scanning, 20: 389–397. 10.1002/sca.1998.4950200504

Zak A, Merino-Cortés SV, Sadoun A, Mustapha F, Babataheri A, Dogniaux S, Dupré-Crochet S, Hudik E, He HT, Barakat AI, Carrasco YR, Hamon Y, Puech PH, Hivroz C, Nüsse O, Husson J. Rapid viscoelastic changes are a hallmark of early leukocyte activation. Biophys J. 2021 May 4;120(9):1692–1704. doi: 10.1016/j.bpj.2021.02.042.

Zucchetti AE, Paillon N, Markova O, Dogniaux S, Hivroz C, Husson J. Influence of external forces on actin-dependent T cell protrusions during immune synapse formation. Biol Cell. 2021 May;113(5):250–263. doi: 10.1111/boc.202000133.

