## Supporting Information for "Stiffening cells with light"

### Stiffening cells with light
#### Supporting Information

Eva Gonzalez[1]*, Jana El Husseiny[2]*, Finn Bastian Molzahn[1], Tiffany Campion[3], Hadrien Jalaber[4], Stéphanie Dogniaux[5], Pierre-Henri Puech[2], Oliver Nüsse[4], Laure Gibot[3] and Julien Husson[1§]

1 Laboratoire d'Hydrodynamique (LadHyX), CNRS, École Polytechnique, Institut Polytechnique de Paris, Palaiseau, France
2 Aix Marseille Université UM61, CNRS UMR 7333, Inserm U1067, Marseille France
3 Laboratoire Softmat, Université de Toulouse, CNRS UMR 5623, Université Toulouse III – Paul Sabatier, 31062 Toulouse, France
4 Institut de Chimie Physique, CNRS UMR 8000, Université Paris-Saclay, Orsay, France
5 Integrative analysis of T cell activation team, Institut Curie-PSL Research University, INSERM U932, Paris, France
* equal contribution

#### Movie Legends

**Movie S1.** Fluo-4 excitation leads to irreversible T-cell deformation and "cellular pancakes". A T cell loaded with Fluo-4 is held by a micropipette and pressed against a rigid wall (a second, larger and filled micropipette) for a few seconds. The holding micropipette is rapidly retracted, and the viscoelastic recovery of the cell shape is monitored. After about 20 s, the cell is pressed a second time against the rigid wall, but while deformed, a blue fluorescence excitation light is turned on (appearance of the star symbol on the movie) and left on. Under the excitation light, the cell does not recover anymore: it appears totally "cooked" (hence informally called a "cell pancake"). A 100x-magnification objective was used. The time is shown in minutes:seconds.

**Movie S2.** Effect on fluorescence excitation during T cell activation. Two Fluo-4-loaded CD4 T cells are activated with anti-CD3+anti-CD28 -coated microbeads. In the absence of fluorescence excitation (above), the T cell emits a large protrusion. In the other cell (below), an exciting blue light is turned on (appearance of the star symbol in the movie) while the protrusion is growing. Very quickly after light application, the protrusion stops growing. A 100x-magnification objective was used. The time is shown in minutes:seconds.

**Movie S3.** Before/after, T cells, DCFH-DA. Profile microindentation of two T cells loaded with DCFH-DA using the "before-after" protocol. Cells are first indented twice (cycle 1-2), and at the end of cycle 2, they are either left in the dark as a control (left) or exposed to 10 s of excitation light (right). After this, they are indented twice to witness the eventual change in stiffness. In these movies, during the 10-s period, the snapshot obtained by accumulating the 10-s exposure is shown as a static image. A 100x-magnification objective was used. The time is shown in minutes:seconds. The time is accelerated 2x.

**Movie S4.** Profile microindentation of a PLB cell loaded with DCFH-DA using the "before-after" protocol. The cell is first indented twice, then exposed to 10 s of excitation light, and then indented again twice to witness the eventual change in stiffness. In this movie no image is shown during the 10-s exposure to blue light. A 100x-magnification objective was used. The time is accelerated 4x.

**Movie S5.** Profile microindentation of four different T cells (columns 1-4) subjected to the "12-cycle two-color". The cells were loaded with both CellTracker Red (CTR) and DCFH-DA. CTR was used as a phototoxic fluorophore, and DCFH-DA was used as a ROS reporter. At the end of each of the 12 indentation cycles, DCFH-DA was excited with 200 ms exposure to blue light at 2% lamp power. From the cycle 3 on, following the short DCFH-DA excitation, a longer 4-s excitation by green light at 100% power was applied to induce CTR-mediated cell stiffening. Images acquired with transmitted light are shown on the first row, and snapshots obtained when exciting CTR are shown below. A 100x-magnification objective was used. The time is shown in minutes:seconds.

**Movie S6.** Timelapse of four T cells loaded with both CellTracker Red (CTR) and DCFH-DA and held by a micropipette (not seeable in the DCFH-DA fluorescence images shown). We performed a 2-color time-lapse microscopy with CTR being used as the phototoxic agent and DCFH-DA as a reporter of ROS production without being too phototoxic. A 200-ms exposure and strong (100% power of the green light) excitation of CTR followed by a shorter 50-ms and low-intensity (10% power of the blue light) excitation of DCFH-DA was performed at a rate of 2 frames per second. Here only the DCFH-DA images are shown. The DCFH-DA fluorescence level increases, reaches a peak and then decreases. A 100x-magnification objective was used. The time is shown in minutes:seconds.

**Movie S7.** T cell loaded with Fluo-4 and submitted to the before-after protocol using a needle to indent the cell. Instead of a single indentation, a compressive force, superimposed with a small modulation, was applied for 10 s to measure the variations in contact stiffness. The movie was acquired at 1 image per second, so the force modulation cannot be seen. A 100x-magnification objective was used. The time is shown in minutes:seconds.

**Movie S8.** Detached endothelial incubated with encapsulated Pheophorbide *a*, submitted to "before-after" protocol but in absence of fluorescence excitation. The time is accelerated 4x.

**Movie S9.** Detached endothelial incubated with encapsulated Pheophorbide *a*, submitted to "before-after" protocol with UV fluorescence excitation. A 100x-magnification objective was used. The time is accelerated 4x.

#### Supplementary Material

| Cell type | condition | Average increase (%) cycle 1 to 2 | N independent experiments | N cells |
|---|---|---|---|---|
| BAEC | DCFH-DA | 16**** | 3 | 34 |
| BAEC | DCFH-DA+DMSO | 37**** | 3 | 44 |
| BAEC | DCFH-DA+LatA/CytoD | 51**** | 3 | 46 |
| BAEC | Empty micelles | 14*** | 2 | 22 |
| BAEC | Control (RPMI) | 11*** | 2 | 15 |
| BAEC | Encapsulated Pheophorbide *a* | 32**** | 3 | 39 |
| BAEC | Pheophorbide *a* | 17** | 4 | 32 |
| BAEC | UV | 11**** | 3 | 20 |
| CD4 | DCFH-DA | 26* | 3 | 34 |
| CD4 | DMSO | 35**** | 3 | 45 |
| CD4 | Hoechst | 29**** | 3 | 30 |
| CD4 | Prolong, Hoechst | 32**** | 2 | 48 |
| PLB | DCFH-DA | 26**** | 5 | 111 |

**Supplementary Table 1**. Increase due to stiffness before fluorescence excitation (comparing stiffness measured during cycle 2 vs. during cycle 1). The cells were never exposed to fluorescence excitation during cycles 1 and 2, so the condition in the second column indicates that the cells were loaded with the mentioned fluorophore, without exciting it with fluorescence (note: these data are thus not exactly those shown in Figure 7C, where cycles 1-2 are average into the “before” value). In the third column, the Mann–Whitney test was used to test if the increase was significant. Labels for p-values are: * for p<0.05, ** for p<0.01, *** for p<0.001, and **** for p <0.0001.

| Cell type | condition | av. incr.(%), cycle 1 to 2 | av. incr.(%), cycle 2 to 3 | *p*-value, 2/1 *vs.* 3/2 | av. incr.( %), cycle 4 to 3 | *p*-value, 4/3 *vs.* 3/2 | *p*-value, 4/3 *vs.* 2/1 | N indep. exper. | N valid cells |
|---|---|---|---|---|---|---|---|---|---|
| BAEC | DCFH-DA no fluor. exc. | 14*** | 6 | 0.0386* | 8 | 0.4307 | 0.0833 | 3 | 16 |
| BAEC | Control (RPMI) | 11*** | 0 | 0.0009*** | 4* | 0.0739 | 0.0315* | 3 | 15 |
| CD4 | DCFH-DA no fluor. exc. | 45* | -10* | 0.0020** | 12* | 0.0342* | 0.0634 | 3 | 17 |
| CD4 | DMSO. no fluor. exc. | 36** | -2 | 0.0020** | 23** | 0.0069** | 0.1354 | 3 | 15 |
| CD4 | Hoechst no fluor. exc. | 25** | 2 | 0.0710 | 13 | 0.1881 | 0.2582 | 3 | 15 |
| PLB | DCFH-DA no fluor. exc. | 32**** | 0 | <0.0001**** | 6 | 0.2134 | 0.0038** | 5 | 55 |

**Supplementary Table 2**. Effect on cell stiffness of a 10-s rest between cycles 2 and 3 without fluorescence excitation. The second column indicates the fluorophore with which the cells were loaded. In these control experiments cells were not exposed to any fluorescence excitation but were left resting in the dark for 10 s between cycles 2 and 3. The Mann–Whitney test was used to test if the increase was significant. Labels for p-values are: * for p<0.05, ** for p<0.01, *** for p<0.001, and **** for p <0.0001.

| Cell type | condition | Average increase (%) cycle 2 to 3 | N independent experiments | N valid cells |
|---|---|---|---|---|
| CD4 | DMSO, blue exc. | 7 | 3 | 15 |
| CD4 | DMSO, no fluor. exc. | -2 | 3 | 15 |
| CD4 | DMSO, UV exc. | 24*** | 3 | 15 |
| BAEC | Control (RPMI) | 0 | 2 | 15 |
| BAEC | UV exc. | 42**** | 3 | 21 |

**Supplementary Table 3**. Effect on cell stiffness of blue and UV light excitation in absence of fluorophore. Labels for p-values are: * for p<0.05, ** for p<0.01, *** for p<0.001, and **** for p <0.0001.

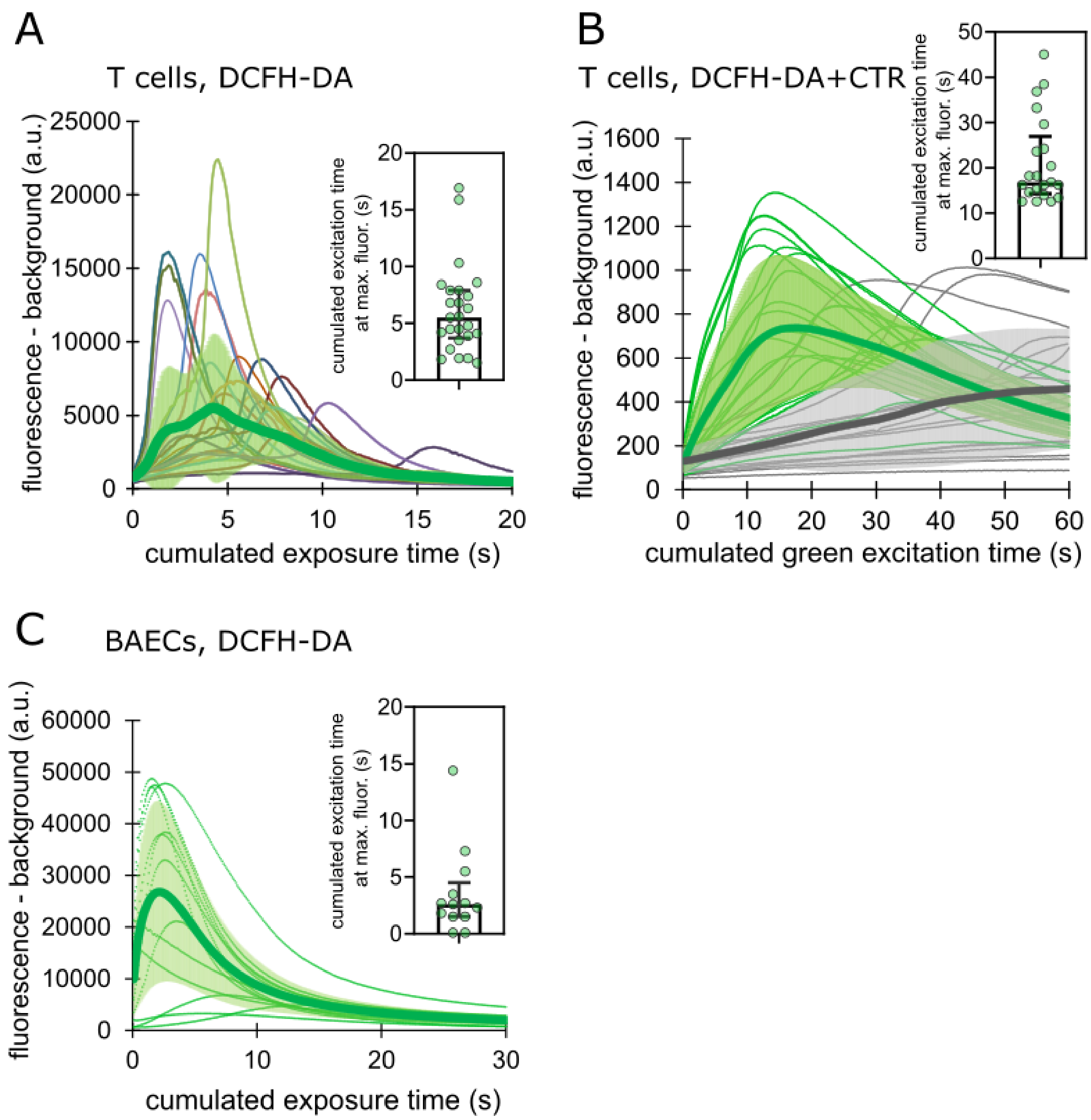


**Suppl. Figure 1.** Level of emitted green fluorescence by DCFH-DA (with subtracted background level) during time-lapse acquisitions. In A-C, thin lines represent single cells, the thick line represents the average, and shaded areas represent standard deviations. **(A)** T cells loaded with DCFH-DA and excited with blue light. Acquisition was performed at one image every 250 ms, exposing cells to blue light at 100% lamp power for 100 ms. Thus, cells were exposed to 400 ms of blue light for every second during the time-lapse, and the x-axis shows the accumulated time of blue-light exposure. Inset: accumulated time of exposure to blue light at which the level of emitted light was maximal. The median time was 5.5 s. Error bars indicate Interquartile ranges. See Movie S6 for four examples of T cells. **(B)** T cells loaded with both CellTracker Red and DCFH-DA and excited with green and blue light. CTR was used as the phototoxic agent, and DCFH-DA at low light doses was used to assess ROS production without excessive phototoxicity. The 2-color time-lapse acquisition was performed with a short exposure (50-ms) and low-intensity (10% power of the blue light) excitation of DCFH-DA and a longer (200-ms) and strong (100% power of the green light) excitation of CTR at a rate of one frame every 500 ms. Cells were exposed to 400 ms of green light for every second during the lapse. The x-axis shows the cumulated time of exposure to green light. Green lines represent cells with green excitation on (i.e. exciting the CTR), and gray lines represent control cells with green excitation off (i.e. not exciting the CTR). Inset: accumulated green-light exposure time at which the level of DCFH-DA emitted light was maximal. The median time was 16.8 s. Error bars indicate interquartile ranges. **(C)** BAECs loaded with DCFH-DA and excited with blue light. Acquisition was performed at one image every 250 ms, exposing cells to blue light at 100% lamp power for 100 ms. Cells were exposed to 400 ms of blue light for every second during the time-lapse, and the x-axis shows the accumulated time of blue-light exposure. Inset: accumulated time of exposure to blue light at which the level of emitted light was maximal. The median time was 2.6 s. Error bars indicate interquartile ranges. See Movie S6 for examples of T cells.

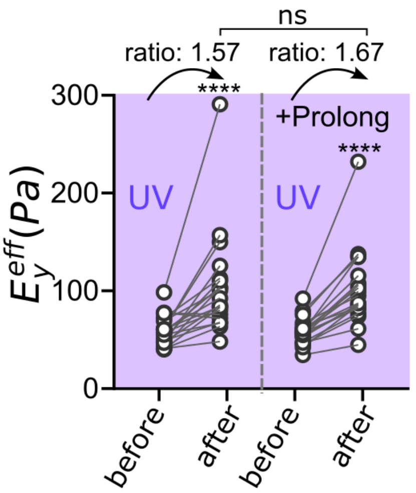


**Supplementary Figure 2.** T cells loaded with Hoechst in the absence or presence of the live antifade reagent Prolong were submitted to the before-after protocol described in Figure 3. T cells become 1.57-fold and 1.67-fold stiffer in the absence and presence of Prolong, respectively. The difference increase ratio is not significant.

#### Supplementary Material 1. Quantification of light intensity on the profile microindentation setup

To measure the irradiance (expressed in mW/cm$^2$) at the specimen level, for a given excitation wavelength of the LED light source, we varied the surface area exposed to excitation light by progressively opening the diaphragm on the path of fluorescent light. We validated that the total power measured by the microscope slide power sensor was proportional to the exposed surface area (Supp. Fig. 3A). The irradiance was taken as the ratio between the measured power and a fixed exposed surface area (3.96x10$^{-5}$ cm$^2$ and 2.49x10$^{-4}$ cm$^2$ for the 100X and 40X objective, respectively). Next, we verified that the power delivered by the lamp increased linearly with the asked power percentage. Although it was not perfectly linear, it could be reasonably approximated as linear (Supp. Fig. 3B). In Supplementary Table 4, the measured irradiance is presented and depends on the combination of objective magnification, excitation wavelength, objective magnification, and filter block.

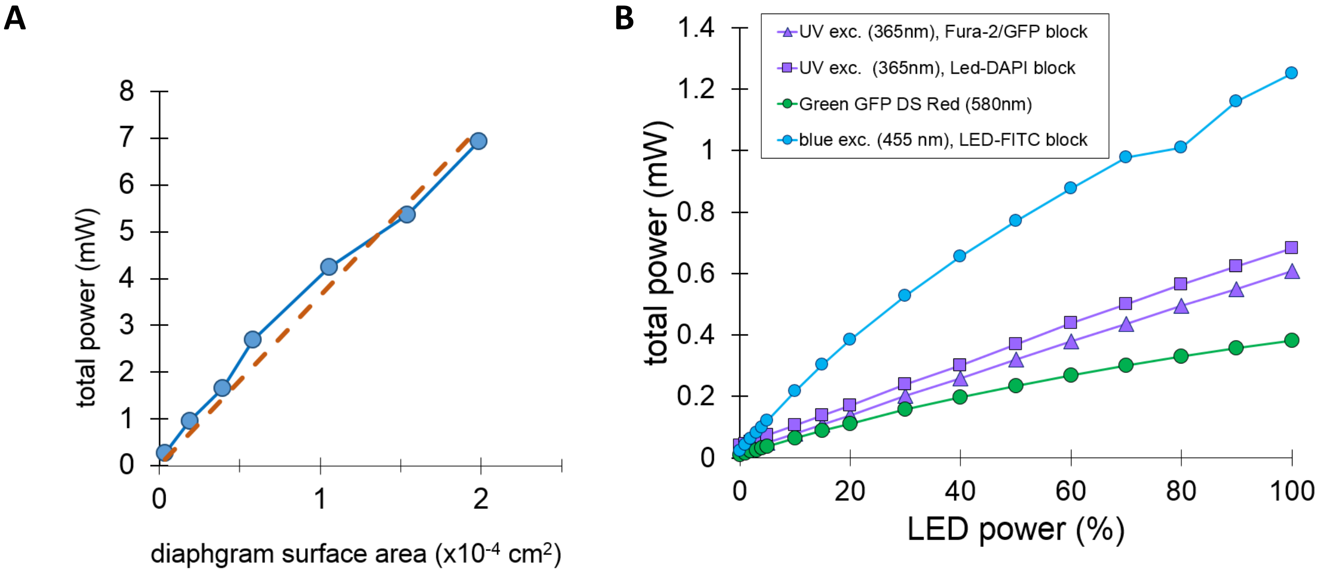


**Supplementary Figure 3.** Light power measured by the microscope slide power sensor on the profile microindentation setup equipped with the 100X-objective. **(A)** Total power as a function of the surface area exposed to excitation light set by opening the diaphragm partially on the path of fluorescent light. **(B)** Total power as a function of LED power at different wavelengths for a fixed exposed area of 3.96x10$^{-5}$ cm$^2$.

| Objective | Fluorophore | block | Excitation wavelength (nm) | Lamp power (%) | Irradiance (mW/cm$^2$) |
|---|---|---|---|---|---|
| 100X | Pheophorbide *a*, Hoechst | LED-DAPI | 365 | 100 | 18 000 |
| | | Fura2-GFP | 365 | 100 | 16000 |
| | DCFH-DA, Fluo-4 | Fura2-GFP | 455 | 100 | 42000 |
| | | LED-FITC | 455 | 100 | 32 000 |
| | CellTracker Red | GFP-DSRED | 580 | 100 | 10000 |
| 40X | DCFH-DA, Fluo-4 | Fura2-GFP | 455 | 100 | 18000 |
| | CellTracker Red | GFP-DSRED | 580 | 100 | 4000 |

**Supplementary Table 4**. Irradiance measured on the profile microindentation setup. irradiance (expressed in mW/cm$^2$) was taken as the ratio between the total power measured by the microscope slide power sensor and a fixed exposed surface area (3.96x10$^{-5}$ cm$^2$ = 3960 µm$^2$ and 2.49x10$^{-4}$ cm$^2$ for the 100X and 40X objective, respectively. Results obtained for different objectives, filter blocks, and excitation wavelengths are presented.

#### Supplementary Material 2. Quantification of light intensity on the AFM setup

The same model of microscope slide power sensor was used to measure the irradiance at the specimen level on the AFM setup. We set the surface area exposed to excitation light to 1600 µm$^2$ by partially opening the diaphragm on the path of fluorescent light. The irradiance was taken as the ratio between the measured power and the exposed surface area (1.59x10$^{-5}$ cm$^2$ = 1590 µm$^2$, using the 40X air objective). We measured 19.7 µW when using 100% blue LED power, so the irradiance was 19.7x10$^{-3}$ mW/1.59x10$^{-5}$ cm$^2$ = 1240 mW/cm$^2$, i.e. 26 to 34 times less irradiance than on the profile microindentation setup using the 100X objective and the LED-FITC or Fura2-GFP filter block, respectively.

#### Supplementary Material 3. Measuring the mechanical properties of RAW macrophages loaded with FLuo-4 or DCFH-DA using atomic force microscopy

During experiments, the AFM cantilever tip was first positioned above the adhered cell using transmitted-light microscopy. The maximal applied force was 500 pN (leading to indentation depths of the order of one µm) using a contact duration of 0 sec. The pressing and pulling speed was 2µm/s, with an imposed maximal displacement of 7 µm. We repeated 5 force curves per cell with no waiting time in-between. Data was typically recorded at 2048 Hz. To determine the Young's modulus of T cells, each experimental force curve was first examined by eye and automatically processed using JPK DP software (JPK Instruments/Bruker). We used the Hertz model for a sphere, and only a subset of the entire force span was fitted, typically over 0.5 µm of indentation, in order to minimize contributions coming from the nucleus, above which we were usually positioned. The Young's modulus measured by analyzing the first force curve, before or after illumination, was considered for a given cell.

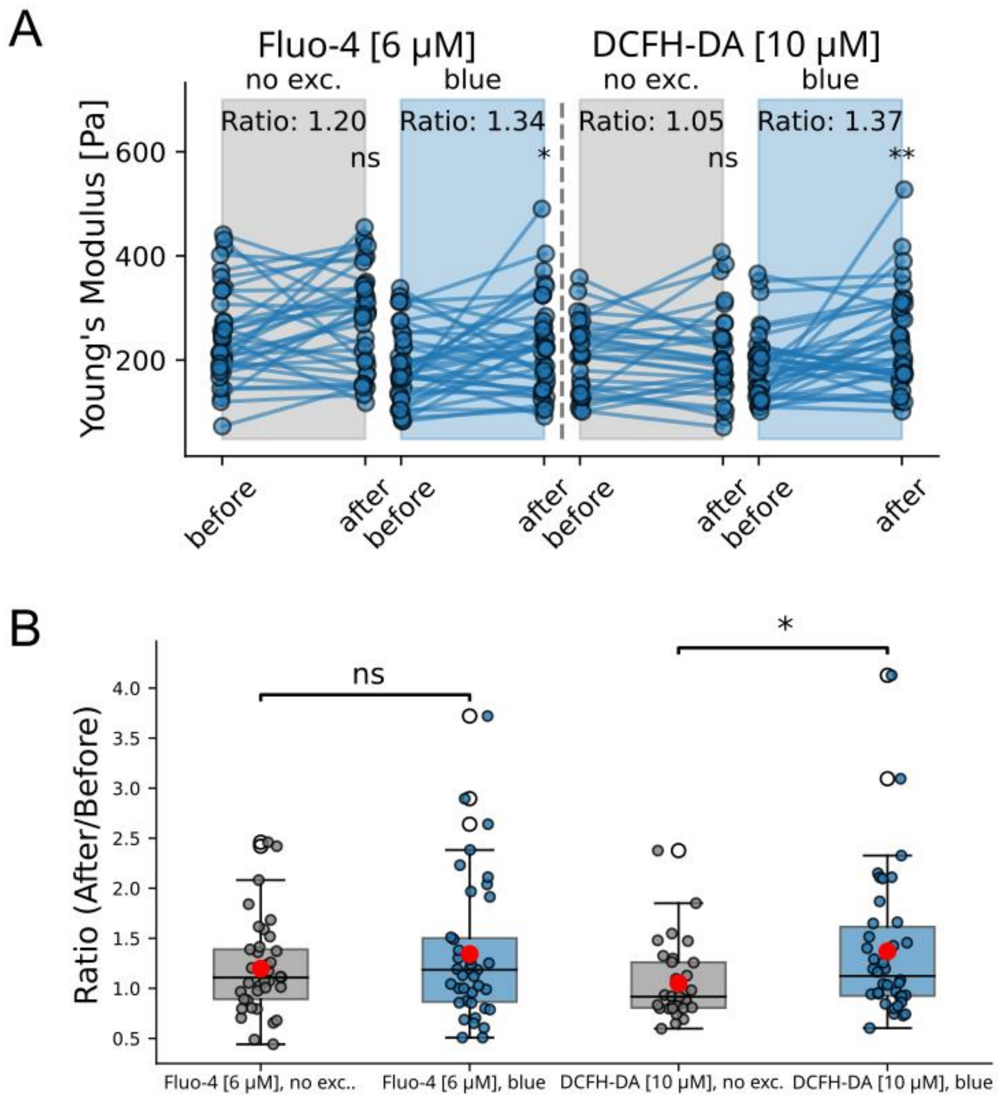


**Supplementary Figure 4.** Photostiffening in RAW cells quantified using AFM. **(A)** Indentation of RAW T cells loaded with Fluo-4 (two columns on the left) or DCFH-DA (two columns on the right) before and after exposure to 30-s blue illumination (blue columns) or without fluorescence excitation (gray columns). **(B)** For the data shown in A, the ratio in the Young's modulus divided by the Young's modulus before fluorescence illumination (or no illumination in control experiments) is shown. The red dot is the average value, the bars and shaded area represent the median and interquartile range. A Mann-Whitney U test on the ratio values shows no significantly higher ratio in Fluo-4 loaded cells, but a significantly larger ratio in DCFH-DA-loaded cells. Statistics: Fluo-4 without fluorescence excitation, n=3 independent experiments, N=37 cells; Fluo-4 with fluorescence excitation, n=3 independent experiments, N=39 cells; DCFH-DA without fluorescence excitation, n=4 independent experiments, N=31 cells; DCFH-DA without fluorescence excitation, n=4 independent experiments, N=38 cells.

#### Supplementary Material 4 Volume rather than cortical stiffening

As explained in detail in [Markova, 2024], indenting a cell with both beads with a diameter of several microns and needles with a tip of submicron radius can help measuring both the cell tension and its inner viscoelastic modulus. In brief, using a sharp tip can probe the cell surface without being influenced by the mechanical properties of the cell interior. As previously done [Markova et al.], we used force modulation to measure the contact stiffness of the needle pressing the surface of a cell. In brief, after a maximal force $F_{max}$ = 100 pN was reached during an initial indentation, we applied a force modulation, i.e., an oscillatory force $F(t) = F_{max} + \Delta F cos(\omega t)$, where $\Delta F$ = 30 pN is the amplitude of the oscillations, $\omega = 2\pi f$ is the angular frequency of the oscillations, and $f$ = 1 Hz is the frequency. Under this oscillatory force, the cell indentation is sinusoidal at the same frequency, $\delta(t) = \delta_m + \Delta\delta cos(\omega t - \varphi)$, where $\delta_m$ is the average of $\delta(t)$, over a period (1 s), $\Delta\delta$ the amplitude of the oscillations (typically 100 nm) and $\varphi$ is a phase lag due to the cell's viscous properties (an applied force leads to a delayed deformation) [Zak 2021]. The amplitude of the force modulation and indentation modulation are related by a complex stiffness defined as $K^* = K' + iK''$, where the part $K' = \frac{\Delta F}{\Delta\delta} cos\ \varphi$ characterizing the elastic properties of the cell and an imaginary part $K'' = \frac{\Delta F}{\Delta\delta} sin\ sin\ \varphi$ characterizing its viscous properties, and $|K^*| = \frac{\Delta F}{\Delta\delta}$ is the norm of the complex stiffness. In this study, for simplicity, we call "contact stiffness" the norm

$|K^*| = \frac{\Delta F}{\Delta \delta}$, which is a model-independent straightforward measurement, because $\Delta F$ is set by the device, and $\Delta\delta$ is directly measured ($|K^*|$ does not depend on the phase lag $\varphi$, which can be more challenging to measure accurately). In previous work, we showed that the contact stiffness for a small contact radius can be can be assumed to be mainly determined by cell tension [Markova, 2024], and is thus a property of the cell surface (its cortex), as opposed to the effective Young's modulus, where the mechanical properties of the cell's interior contribute as well.
We asked whether photostiffening mainly affected the cell interior and not its surface. If so, we expected that the contact stiffness would not increase, whereas the effective Young's modulus would. Thus, we performed indentation with both beads and needles. As shown in Figure 7, in T cells loaded with Fluo-4, the stiffness increased 2.6-fold after indenting with a sphere (Fig 7A), which was significantly more than the 1.5-fold increase in contact stiffness measured after indenting with a needle (Fig 7B).

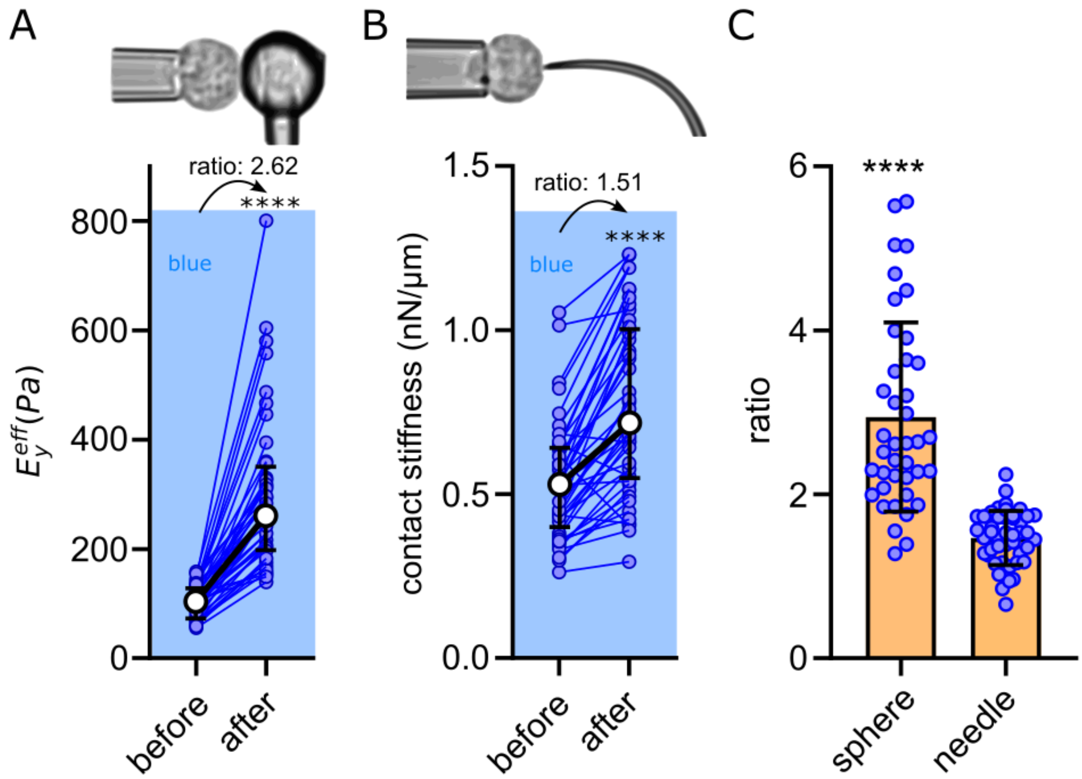


**Supplementary Figure 5. (A)** Stiffness of T cells loaded with Fluo-4 and submitted to the before-after protocol. The cell stiffness increased by a factor of 2.62. Data are reproduced from Figure 5A. **(B)** Instead of a spherical indenter, a needle was used to indent cells, and instead of a single indentation, a modulated compressive force was applied for 10 s to measure the contact stiffness of Fluo-4-loaded T cells (see Materials and Methods and Movie S7). The contact stiffness increased by a factor of 1.51 after fluorescence excitation. **(C)** The ratio of increase in effective Young's modulus measured with a special indenter is significantly larger than the increase in contact stiffness measured using a needle.
